# The *Medicago truncatula* DREPP protein triggers microtubule fragmentation in membrane nanodomains during symbiotic infections

**DOI:** 10.1101/794008

**Authors:** Chao Su, Marie-Luise Klein, Casandra Hernández-Reyes, Morgane Batzenschlager, Franck Anicet Ditengou, Jean Keller, Pierre-Marc Delaux, Thomas Ott

## Abstract

The initiation of intracellular host cell colonization by symbiotic rhizobia in *Medicago truncatula* requires repolarization of root hairs, which includes the re-arrangement of cytoskeletal filaments. The molecular players governing microtubule (MT) re-organization during infection remain to be discovered. Here, we identified the *M. truncatula* DREPP protein and investigated its functions during rhizobial infections. We show that rhizobial colonization of *drepp* mutant roots as well as transgenic roots over-expressing DREPP is impaired. DREPP re-localizes into symbiosis-specific membrane nanodomains in a stimulus-dependent manner. This subcellular segregation coincides with DREPP-dependent MT fragmentation and a partial loss of the ability to re-organize the MT cytoskeleton in response to rhizobia, which might relay on an interaction between DREPP and MT organizing protein SPIRAL2 (SPR2). Taken together, our results reveal that establishment of symbiotic associations in *M. truncatula* require DREPP in order to regulate MT reorganization during initial root hair responses to rhizobia.

## Introduction

The legume-Rhizobium root nodule symbiosis (RNS) represents an almost exceptional example for an intracellular accommodation of a bacterial mutualist. In the model legume *Medicago truncatula* rhizobial infection strictly depends on the molecular recognition of strain-specific rhizobial signalling molecules, called Nod Factors (NFs) (Oldroyd, 2013). These are secreted by its symbiont *Sinorhizobium meliloti* and are perceived by the host LysM-type receptor-like kinases (RLKs) NOD FACTOR PERCEPTION (NFP) (Amor et al., 2003; Arrighi et al., 2006) that possibly form a complex with the entry receptor LYSIN MOTIF RECEPTOR KINASE 3 (LYK3) (Limpens et al., 2003; Smit et al., 2007). This pre-infection molecular dialogue also requires the function of DOES NOT MAKE INFECTIONS 2 (DMI2) (Catoira et al., 2000; Stracke et al., 2002; Hogg et al., 2006), an LRR-RLK that acts in concert with NFP and LYK3 (Antolin-Llovera et al., 2014). All three receptors physically interact with the molecular scaffold protein SYMREM1 (Lefebvre et al., 2010; Toth et al., 2012; Liang et al., 2018) in membrane nanodomains. Localizations of RLKs to such sub-micron protein/lipid assemblies have also been described for other receptors such as the innate immune receptor FLAGELLIN SENSING 2 (FLS2) and the brassinosteroid receptor BRI1 (Bucherl et al., 2017). Since membrane nanodomains also host other signaling components, they have been hypothesized to play a major role for subcompartmentalizing the PM into functionally distinct (reviewed in Ott, 2017; Yu et al., 2019). While the LYK3 receptor displays a comparably high lateral mobility at the PM prior to infection (Haney et al., 2011) it gets immobilized in membrane nanodomains labelled by FLOTILLIN 4 following infection by rhizobia (FLOT4; Haney and Long, 2010; Haney et al., 2011). This immobilization is mediated by the remorin protein SYMREM1 and results in a stabilization of LYK3 during infection (Liang et al., 2018). Genetic evidence suggests that LYK3 recruitment into FLOT4/SYMREM1-positive nanodomains is required for successful rhizobial infections (Liang et al., 2018).

Ligand-dependent activation of the primary symbiotic receptors NFP, LYK3 and DMI2 ultimately results in regular nuclear and perinuclear calcium oscillations, a hallmark for symbiotic infections (Ehrhardt et al., 1996; Kosuta et al., 2008). These calcium spikes are decoded by a calcium-calmodulin-dependent kinase (DMI3) (Levy et al., 2004), which in turn activates, among others, the transcription factor CYCLOPS/IPD3 via phosphorylation (Messinese et al., 2007; Yano et al., 2008; Hayashi et al., 2010; Singh et al., 2014). As a consequence, two distinct genetic programs are triggered: one controlling different steps of rhizobial infection and the other one the organogenetic pathway that results in the formation of root nodules (Madsen et al., 2010).

In *M. truncatula*, infection requires a transient growth retardation of susceptible root hairs that results in root hair swelling and a subsequent deformation. This redirects the resumed root hair growth and leads to the formation of a root hair curl (RHC) that entraps a single rhizobial colony within the so-called infection chamber (IC) (Fournier et al., 2015). Upon rhizobial divisions inside the IC, the plasma membrane of the host invaginates, leading to the formation of a tunnel-like structure, the infection thread (IT) that continues growing into the newly formed nodule primordium (Gage, 2004; Fournier et al., 2008; Fournier et al., 2015). Here, rhizobia will be released into infected cells, where they differentiate into nitrogen-fixing bacteroids (Gage, 2004; Montiel et al., 2017). In so called ‘indeterminate nodules’, as formed by *M. truncatula*, these infected cells undergo senescence at the base of the nodule (zone IV) while newly infected cells are formed close to the nodule apex zone II (Vasse et al., 1990).

The initial root hair responses are probably mediated by F-actin that undergoes significant re-arrangements enabling the induction of root hair deformation, root hair curling and infection thread formation (Crdenas et al., 1998; Yokota et al., 2009; Hossain et al., 2012; Zepeda et al., 2014). Later on, a polarity switch is required to re-direct polar secretion towards the IC (Fournier et al., 2015) and IT growth (Fournier et al., 2008). Among other genetic components the *Lotus japonicus* LjSCARN (SCAR-Nodulation), an activator of the ARP2/3 actin nucleation complex, is required during root hair polar growth (Qiu et al., 2015). Similar to mutations in other members of the SCAR/WAVE complex, such as NAP1 and PIR1 (Yokota et al., 2009), *scarn* mutants display short root hairs and mostly transverse or web-like actin filaments compared to WT with a predominantly longitudinally oriented actin cytoskeleton (Qiu et al., 2015). Further involvement of the ARP2/3 complex at later stages of the symbiotic interaction was presented in *M. truncatula*, where MtARP3 controls symbiosome development (Gavrin et al., 2015). By contrast, our knowledge on the molecular regulation of microtubule (MT) arrangements during symbiotic interactions is comparably sparse. It has been shown that only those root hairs that just stopped elongating are able to respond to NFs and rhizobia by root hair curling (Heidstra et al., 1994). Here, MT growth rates are decreased by around 45% suggesting an active role in this process (Vassileva et al., 2005). Additionally, imaging of Rhizobium-treated root hairs revealed that the reorganization of MTs is tightly linked to nuclear repositioning during root hair curling and IT initiation (Timmers et al., 1999). Despite the unequivocal importance of MT patterning during rhizobial infections, these processes have not been genetically dissected. Interestingly, plant-specific DEVELOPMENTALLY-REGULATED PLASMA MEMBRANE POLYPEPTIDE (DREPP) proteins were shown in *A. thaliana* to regulate the actin and MT cytoskeleton in a calcium-dependent manner (Li et al., 2011; Zhu et al., 2013; Qin et al., 2014). The DREPP protein family is comprised of only two members in *A. thaliana*: The PLASMA MEMBRANE-ASSOCIATED CATION-BINDING PROTEINs PCaP1 (AtMDP25) and PCaP2 (AtMAP18) (Li et al., 2011; Zhu et al., 2013; Qin et al., 2014). PCaP1 is sufficient to sever MTs and actin filaments in vitro, while this function has not been confirmed in vivo. The process is enhanced upon calcium treatment and additionally results in the dissociation of PCaP1 from the membrane to the cytosol. In the presence of calcium PCaP2 interacts with calmodulin (CaM) and phosphatidylinositol phosphates (PtdInsPs) (Nagasaki et al., 2008).

Here, we demonstrate that *M. truncatula* DREPP is required for successful infection by rhizobia. The application of calcium and Nod Factors resulted in a re-localization of DREPP into nanodomains that co-localize with MTs. This accumulation coincides with fragmentation of MT in root hairs. Given the ability of DREPP to interact with MTs and MT-associated proteins, we propose that DREPP processes MTs in functional membrane nanodomains.

## Results

### *drepp* mutants are affected in rhizobial infections and nodulation

Rhizobial infections in legumes are accompanied by significant cytoskeleton re-arrangements prior to and during infection thread progression. To understand this phenomenon in more detail, we studied the single member (Medtr2g437530) of the DREPP protein family in *M. truncatula*. Large-scale phylogenetic analysis using 104 plant genomes revealed an unambiguous presence of DREPP paralogs in 91 species, indicating that DREPP is conserved throughout the angiosperms, i.e. in species that are able to nodulate but also species from lineages that have lost, or never evolved nodulation (Supplemental Figure S1; Table S1). In *M. truncatula DREPP* transcripts can be continuously detected and are transiently but mildly induced about three days post inoculation (dpi) of roots with rhizobia (Supplemental Figure S2A). Fusing a 1.5kb long fragment of the putative *DREPP* promoter to β-glucuronidase (GUS) revealed promoter activity in root cortical cells and root hairs at 7 dpi (Supplemental Figure S2B). Residual promoter activity of *DREPP* was also observed in the two receptor mutants *nfp* and *dmi2* as well as in the *dmi3* mutant background (Supplemental Figure S2B) indicating that basal expression is, as expected, independent of the induction of symbiotic signaling.

To genetically asses the relevance of DREPP during root nodule symbiosis, we isolated and analyzed two independent *drepp* mutant alleles, *drepp-1* (NF1952) and *drepp-2* (NF8505), carrying TNT1 transposon insertions at positions 1038bp and 988bp downstream of the translational start codon of the *DREPP* gene, respectively (Fig. 1A). Quantitative RealTime PCR revealed that endogenous *DREPP* transcripts were significantly decreased by about 10-fold in homozygous *drepp-1* and *drepp-2* mutant plants confirming that they can be considered as transcriptional knockdown alleles (Fig. 1B). Since DREPP proteins have been reported to function during root hair (RH) development and nuclear movement in root hairs (Zhang et al., 2015), which would indirectly interfere with symbiotic infections, we inspected RH development and additionally scored the distance between the nucleus and the tip of elongating root hairs in the susceptible zone around 10 mm above the root cap. No significant differences were observed between the genotypes (Supplemental Figure S3). To confirm, however, that DREPP serves functions in MT re-arrangements as described for the orthologous protein from *A. thaliana* (Wang et al., 2007), we performed immunofluorescence using an anti-tubulin antibody to visualize cortical MT orientation in wild type (WT, R108), *drepp* mutants and transgenic roots constitutively over-expressing *DREPP* (DREPP-OE). Parallel arrays of transversely oriented MT were predominantly observed in WT root epidermal cells (Supplemental Figure S4). While *drepp* mutants displayed mostly transversely oriented MTs and only few with random MT arrays (Supplemental Figure 4), ectopic expression of *DREPP* resulted in a significant re-orientation of MTs with random, oblique, or longitudinal MT alignments in root epidermal cells (Supplemental Figure S4). This indicates that the *M. truncatula* DREPP protein retained conserved but may have lost pleiotropic functions.

**Fig. 1.**
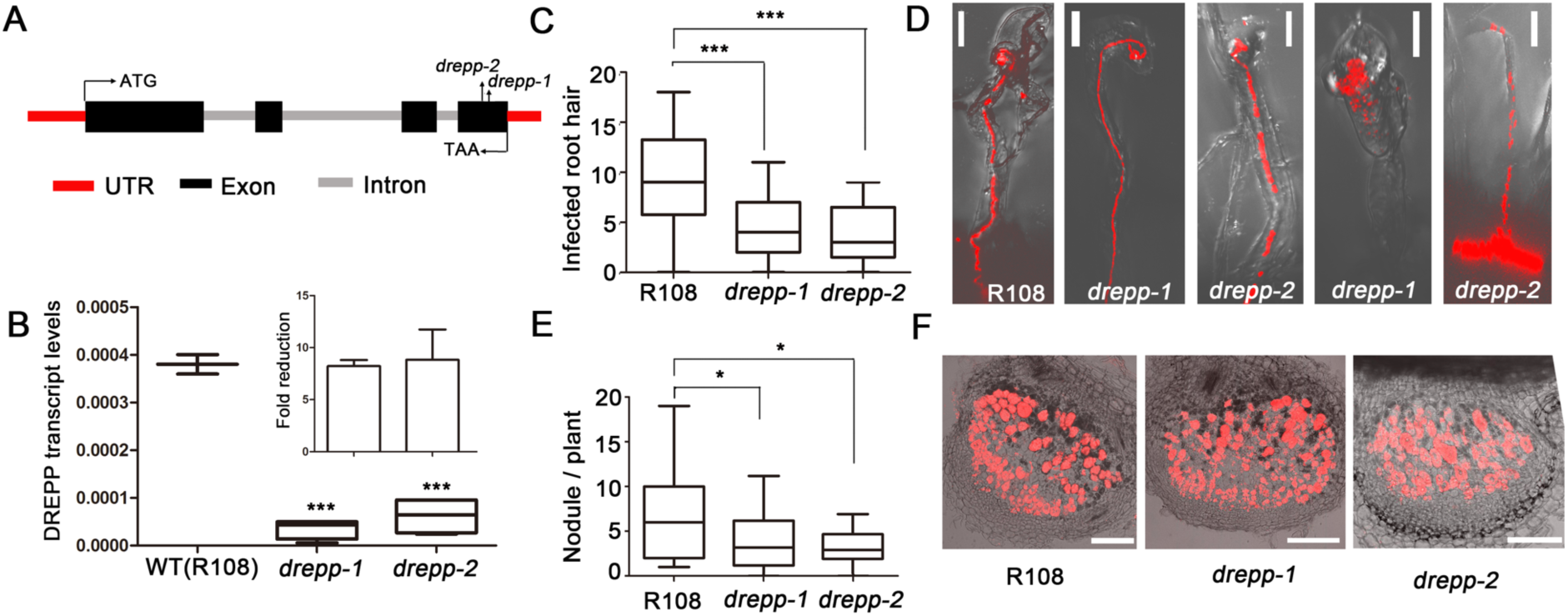
DREPP regulates rhizobial infections. (**A**) Schematic representation of the *DREPP* gene structure and mapped *Tnt1* transposon insertion sites in two independent *drepp* mutant alleles. UTR= untranslated region. (**B**) Endogenous transcript levels were quantified by qRT-PCR. Values are normalized to the housekeeping gene *Ubiquitin.* Infected root hairs (curled root hairs) (**C**) and nodules (**E**) were scored at 7 dpi with n ≥19 root systems for each genotype and condition. (**D**) While several root hairs developed WT-like ITs in *drepp-1* and *drepp-2* mutants, a great proportion was pre-maturely aborted or bacteria were released into trichoblasts (two most right panels), Scale bars indicate 20 µm. (**F**) Semi-thin (70µm) longitudinal sections of 7 dpi old whole nodules from wild-type and *drepp* mutants. Red fluorescence derives from mCherry-labeled *S. meliloti* hosted in infected nodule cells, Scale bars indicate 100 µm. Data are given as mean ± SE. Statistics were performed by using an unpaired, two-tailed Student t test (and nonparametric tests). *= p<0.05; ***= p< 0.001.

**Fig. 2.**
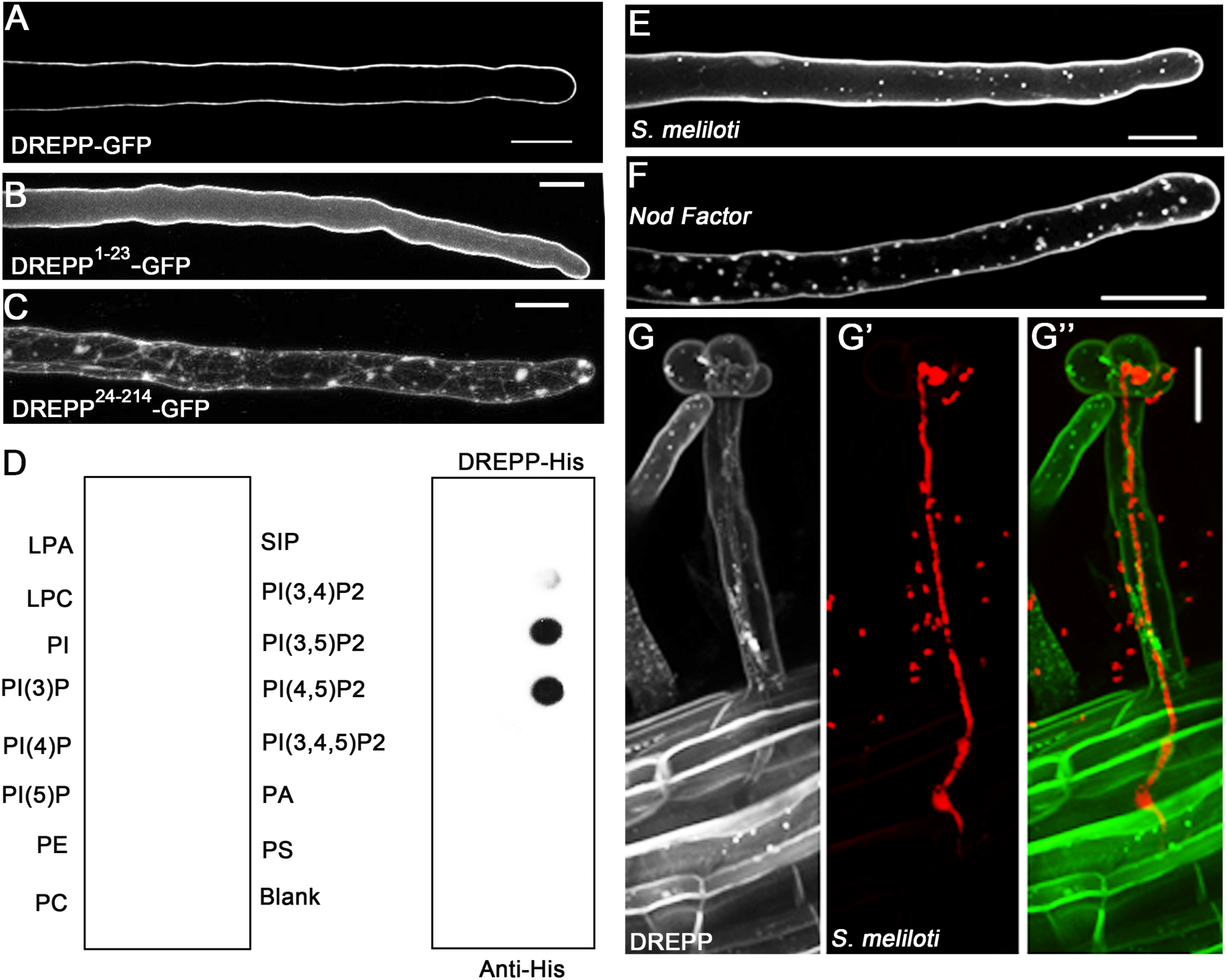
DREPP binds to the PM and relocalizes to membrane nanodomains in a symbiosis-dependent manner. (**A**) A DREPP-GFP fusion protein localizes to the PM in uninoculated *M. truncatula* root hairs. (**B**) The N-terminal 23 amino acids (1-23) are required (**C**) and sufficient (**B**) to associated soluble GFP with the PM. (**D**) Hybridization of recombinant His-tagged DREPP to PIP Strips™ revealed a specific association with PI(3,4)P2, PI(3,5)P2 and PI(4,5)P2. (**E-F**) Inoculation and application of DREPP-GFP expressing root hairs with *S. meliloti* for 7 days (**E**) and isolated NFs for 12 hours (**F**) results in a specific accumulation of DREPP in membrane nanodomains as indicated by punctate structures. (**G-G”**) Nanodomain localization of DREPP is abolished in IT-containing root hairs (harboring mCherry-labelled rhizobia) but present in cortical cells prior to infection. Scale bars indicate 20 µm.

**Fig. 3.**
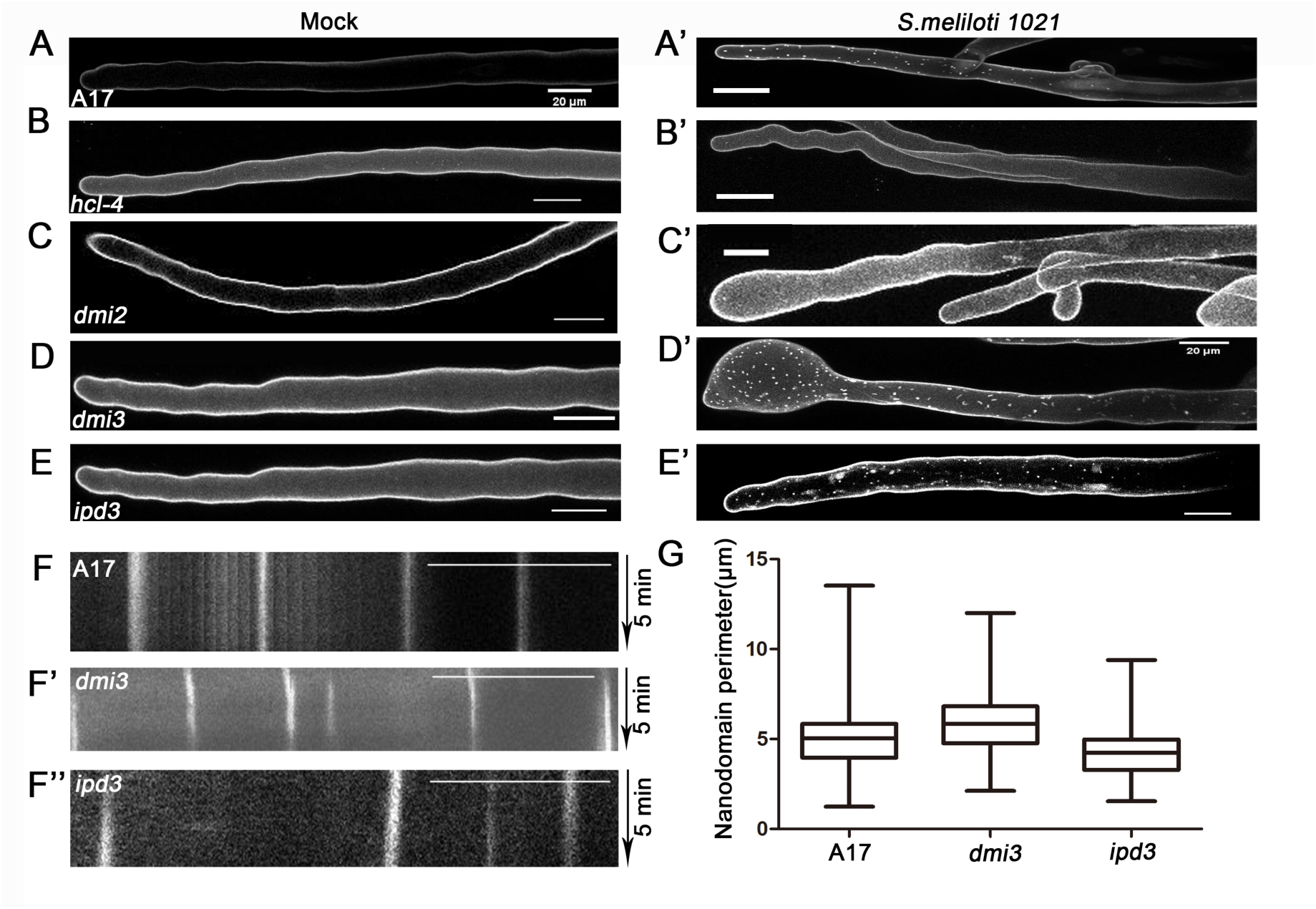
Genetic requirements for nanodomain recruitment of DREPP. (**A-E**) DREPP-GFP localizes homogenously to the PM of root hairs in different Medicago mutants in the absence of rhizobia. (**A’-E’**) Relocalization of DREPP to membrane nanodomains occurred in a symbiosis-dependent manner at 7 dpi in all genotypes except in the receptor mutants *hcl-4* (**B’**) and *dmi2* (**C’**). (**F-F”**) Kymograph analysis of an ∼80 frame time-lapse (5 min) image series to determine spatio-temporal mobility of nanodomain-localized DREPP in WT (A17; **F**), *dmi3* (**F’**) and *ipd3* (**F”**). Vertical lines indicate full immobility of the protein. (**G**) ImageJ-based analysis of nanodomain perimeters did not reveal any significant difference between the genotypes. Data are given as mean ± SE. Statistics were performed by using an unpaired, two-tailed Student t-test (and nonparametric tests).

**Fig. 4.**
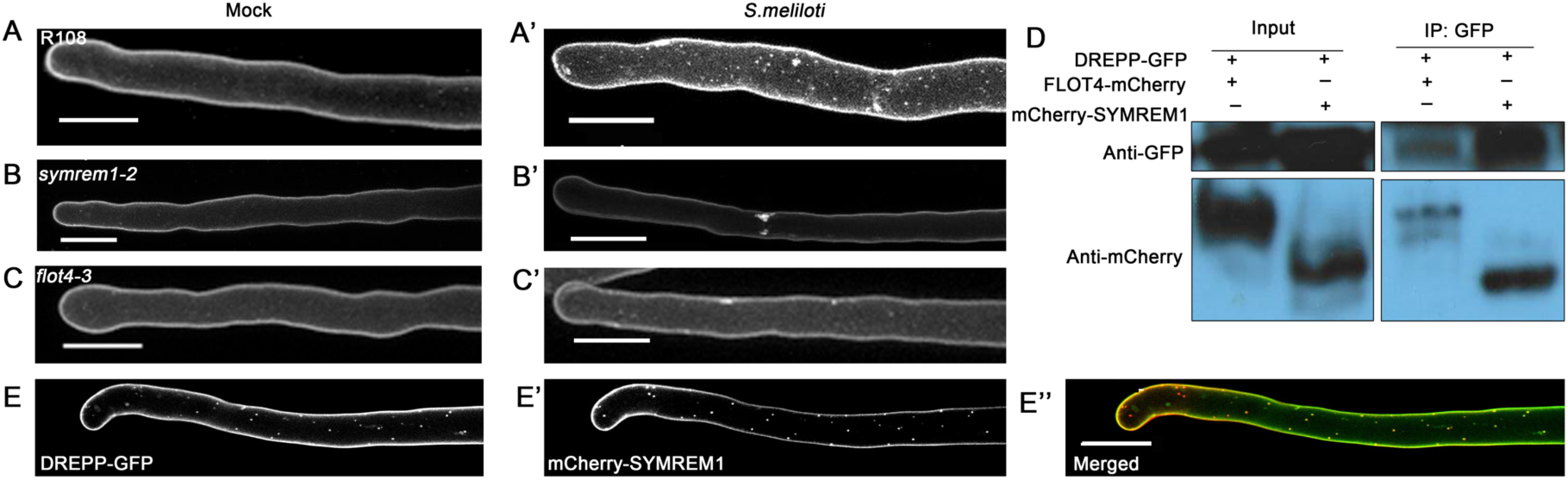
DREPP localizes to membrane nanodomains in a FLOT4- and SYMREM1-dependent manner. (**A-C’**) *M. truncatula* root hairs transformed with *pUBI:DREPP:GFP* in WT (R108) (**A-A’**), *symrem1-2* (**B-B’**) and *flot4-3* (**C-C’**) under non-inoculated (mock) conditions (**A-C**) and after 7 dpi with *S. meliloti* (**A’-C’**). Images are maximum intensity z-projections of ∼70 optical sections acquired at 0.5-1 µm increments. (**D**) Immuno-precipitation experiment after pairwise co-expression of DREPP with SYMREM1 or FLOT4 in *N. benthamiana* leaves using anti-GFP nanotraps revealed DREPP to be a component of the FLOT4-SYMREM1 complex. (**E-E”**) Co-expression and -localization of DREPP-GFP (**E**) and mCherry-SYMREM1 (**E’**) in transgenic *M. truncatula* A17 root hairs at 7dpi with *S. meliloti*. Scale bars indicate 20 µm.

### DREPP regulates rhizobial infections

To investigate whether DREPP is important for rhizobial infections, we phenotypically characterized *drepp* mutants by assessing primary infection success of rhizobia at 7 dpi. For this, we scored responsive root hairs (curled root hairs with or without infection threads (ITs)) and nodule numbers. The number of responsive root hairs was significantly reduced in both *drepp* mutants compared to WT plants (Fig. 1C) even though several mature ITs were observed (Fig. 1D). As a consequence, *drepp* mutants developed less nodule primordia at 7 dpi (Fig. 1E), which were fully colonized by rhizobia and did not exhibit any differences compared to the WT (Fig. 1F). Interestingly, the infection patterns of roots over-expressing DREPP resembled the mutant phenotype (Supplemental Figure S5). Furthermore, we systematically found young elongated nodules with impaired bacterial release (Supplemental Figure S6). In addition, 4-weeks old *drepp* mutant nodules showed an extended nodular senescence zone (zone IV) at their base (Supplemental Figure S6).

### DREPP localizes to membrane nanodomains in response to symbiotic stimuli

To elucidate the molecular function of DREPP during symbiotic infections we first examined the localization of a DREPP-GFP fusion protein. Ectopic expression of *DREPP-GFP* resulted in a peripheral fluorescent signal in young root hairs, indicating an association of the protein with the plasma membrane (PM) (Fig. 2A). PM localization is mediated by the N-terminal region of the protein since the N-terminal 23 amino acids (DREPP^1-23^) of the DREPP polypeptide were required and sufficient to immobilize GFP at the PM (Fig. 2B). Reciprocally, deletion of the N-terminal 23 amino acids (DREPP^24-214^) resulted in a cytosolic accumulation of the protein (Fig. 2C). As the DREPP polypeptide harbors a glycine residue after the N-terminal methionine, we tested whether this residue serves as a myristoylation site by expressing a mutant variant where we exchanged the glycine by an alanine (DREPP^G2A^) in *Nicotiana benthamiana* leaf epidermal cells. While the wild-type DREPP protein localized to the PM, the DREPP^G2A^ mutant showed an increased cytosolic fraction although the majority of the protein remained at the PM (Supplemental Figure S7A). A stronger effect was observed when generating an N-terminal GFP fusion with DREPP (GFP-DREPP), where the presence of GFP abolishes the recognition of the glycine residue as a myristoylation site in the DREPP polypeptide (Supplemental Figure S7A). Due to a strongly increased cytosolic fraction of the GFP-DREPP protein (Supplemental Figure S7B, lane 6) compared to the DREPP^G2A^ mutant we assumed an additional mechanism to contribute to PM binding. Since PM association of tobacco DREPP2 is mediated by a strong polybasic cluster in its N-terminal domain (Vosolsobe et al., 2017), we predicted surface exposed residues in the *M. truncatula* DREPP protein. Indeed, four lysine residues (K7, K11, K14, K18) were found to form a putative polybasic cluster (Supplemental Figure 7C). To test the impact of this poly-lysine stretch on PM association experimentally, we generated three mutant variants where we replaced one (DREPP^K14E^, DREPP^K18E^) or two lysines (DREPP^K14/18E^) by acidic glutamates. While expressing DREPP^K14E^ and DREPP^K18E^ resulted in more visible cytosolic strands (Supplemental Figure S7A) and an increased cytosolic fraction (Supplemental Figure S7B), this effect was further pronounced in the DREPP^K14/18E^ double mutant (Supplemental Figure S7). As polybasic clusters have been reported to support the association of proteins with phosphoinositolphosphates (PIPs), we recombinantly expressed and purified full-length Medicago DREPP from *E. coli* and hybridized it to PIP-strips containing a series of different PIPs. Indeed, DREPP strongly bound to PI(3,5)P_2_ and PI(4,5)P_2_ (Fig. 2D). Taken together these data suggest a bifunctional mechanism of PM association for DREPP.

Since *A. thaliana* PCaP1 was shown to function in the presence of calcium (Li et al., 2011; Zhu et al., 2013; Qin et al., 2014), we tested whether the application of calcium alters PM localization of the DREPP-GFP fusion protein. Expression of DREPP-GFP under the control of the native DREPP promoter (1.5 kb) resulted in a weak but clear signal at the PM of root hairs (Supplemental Figure S8A). Interestingly, application of 1mM CaCl_2_ for two hours induced a recruitment of the protein into membrane nanodomains (Supplemental Figure S8A). The same effect was observed when ectopically expressing DREPP using the *L. japonicus* Ubiquitin10 promoter (Supplemental Figure S8A). Due to the low fluorescence of the pDREPP:DREPP-GFP construct, we continued using the ubiquitin promoter driven construct. To study the dynamics of DREPP during nodulation, we inoculated roots constitutively expressing DREPP-GFP with symbiotic *Sinorhizobium meliloti* (Fig. 2E) or isolated *S. meliloti* Nod Factors (NFs) (Fig. 2F). Here, we observed a re-localization of DREPP into membrane nanodomains at 7dpi and 12 hpi, respectively. However, these structures almost fully disappeared in IT-containing root hairs but were present in cortical cells prior to their penetration by the IT (Fig. 2G).

To genetically position the recruitment of DREPP into these membrane nanodomains, we expressed DREPP-GFP in different mutant backgrounds that show symbiotic phenotypes. In the absence of rhizobia, a peripheral GFP signal was observed, indicating that DREPP resided in the PM in all tested genotypes (Fig. 3A-E). Interestingly and different to the WT (A17; Fig. 3A’) the lateral segregation of DREPP into nanodomains was abolished in the LYK3 and DMI2 receptor mutant alleles *hcl-4* and *dmi2*, respectively (Fig. 3B’-C’). By contrast, DREPP localization was unaffected in *dmi3* and *ipd3* mutants (Fig. 3D-E’). Interestingly, nanodomain recruitment of DREPP was rescued in *dmi2, hcl-1* and *hcl-4* mutants by exogenous calcium application (Supplemental Figure S8B). As in A17 wild-type root hairs, membrane nanodomains were laterally stable over 5 minutes (Fig. 3F-F”), which is a hallmark of these membrane subcompartments in plants (Jarsch et al., 2014; Liang et al., 2018). To address if DREPP is recruited into a recently described symbiosis-induced nanodomain (Haney et al., 2011; Liang et al., 2018), we tested whether DREPP localization depends on the two nanodomain scaffolds SYMREM1 and FLOT4 by expressing DREPP in the *symrem1-2* and the *flot4-3* mutant backgrounds. While DREPP uniformly localized to the PM in the absence of rhizobia (Fig. 4A-C), nanodomain recruitment was impaired in both mutant alleles (Fig. 4B’-C’) but not in the corresponding R108 wild-type (Fig. 4A’). Furthermore, calcium treatment was not sufficient to induce nanodomain recruitment of DREPP in *flot4* and *symrem1* mutants (Supplemental Figure S8B). These data implied that FLOT4 and/or SYMREM1 are essential to recruit DREPP into membrane nanodomains. To test whether both scaffolds are able to associate with DREPP, we pairwise co-expressed *DREPP* with *FLOT4* or *SYMREM1* in *N. benthamiana* leaf epidermal cells and performed co-immunoprecipitation experiments by precipitating GFP-tagged DREPP using anti-GFP nanotraps. Indeed, DREPP co-purified with both FLOT4 and SYMREM1 indicating that these proteins form a complex (Fig. 4D). Since FLOT4 and SYMREM1 co-localize in the same nanodomain upon rhizobial inoculation (Liang et al., 2018), we hypothesized that DREPP should be recruited into the same SYMREM1-positive structures. To test this, we co-expressed *DREPP* and *SYMREM1* in *M. truncatula* roots and inoculated these transgenic roots for 7 days with *S. meliloti*. Both proteins perfectly co-localized in immobile nanodomains under these conditions (Fig. 4E-E”) demonstrating that DREPP is recruited to this symbiosis-induced nanodomain in a stimulus-dependent manner.

### DREPP nanodomains are linked with microtubule fragmentation

Since few DREPP proteins have been shown to function in MT processing in vitro, we first tested a possible association of DREPP with MTs using an in vitro co-sedimentation assay. For this, we used purified dimers of α- and β-tubulin and made use of their self-assembling capacity at 30°C in the presence of GTP and paclitaxel, a MT stabilizing agent. When adding recombinantly expressed and purified the His-tagged DREPP protein, it associated with polymerized MTs directly as indicated by the presence of DREPP in the MT pellet (Supplemental Figure S9). Next, we asked whether MT association and possible processing occurs in membrane nanodomains. For this, we co-expressed *DREPP-mCherry* and the MT marker *YFP-MAP4* in *M. truncatula* root hairs. As shown before in the root epidermis, ectopic expression of *DREPP* resulted in a reorientation of MTs but the filaments appeared largely intact (Supplemental Figure 4). This pattern was also observed in root hairs (Fig. 5). However, when scoring DREPP localization and MT fragmentation after inoculating roots for 7 days with *S. meliloti*, i.e. after DREPP re-localization into nanodomains, MTs were found to be partially fragmented in wild-type roots but not in the *drepp* mutants (Fig. 5). This effect was largely pronounced when over-expressing DREPP (Fig. 5). Here, DREPP-labelled nanodomains aligned with MTs (Supplemental Figure S10). However, treating uninoculated root hairs with the MT depolymerizing drug oryzalin did not prevent recruitment of DREPP to the PM (Supplemental Figure S11) indicating that DREPP processes MTs at the PM rather than requires them for its localization.

**Fig. 5.**
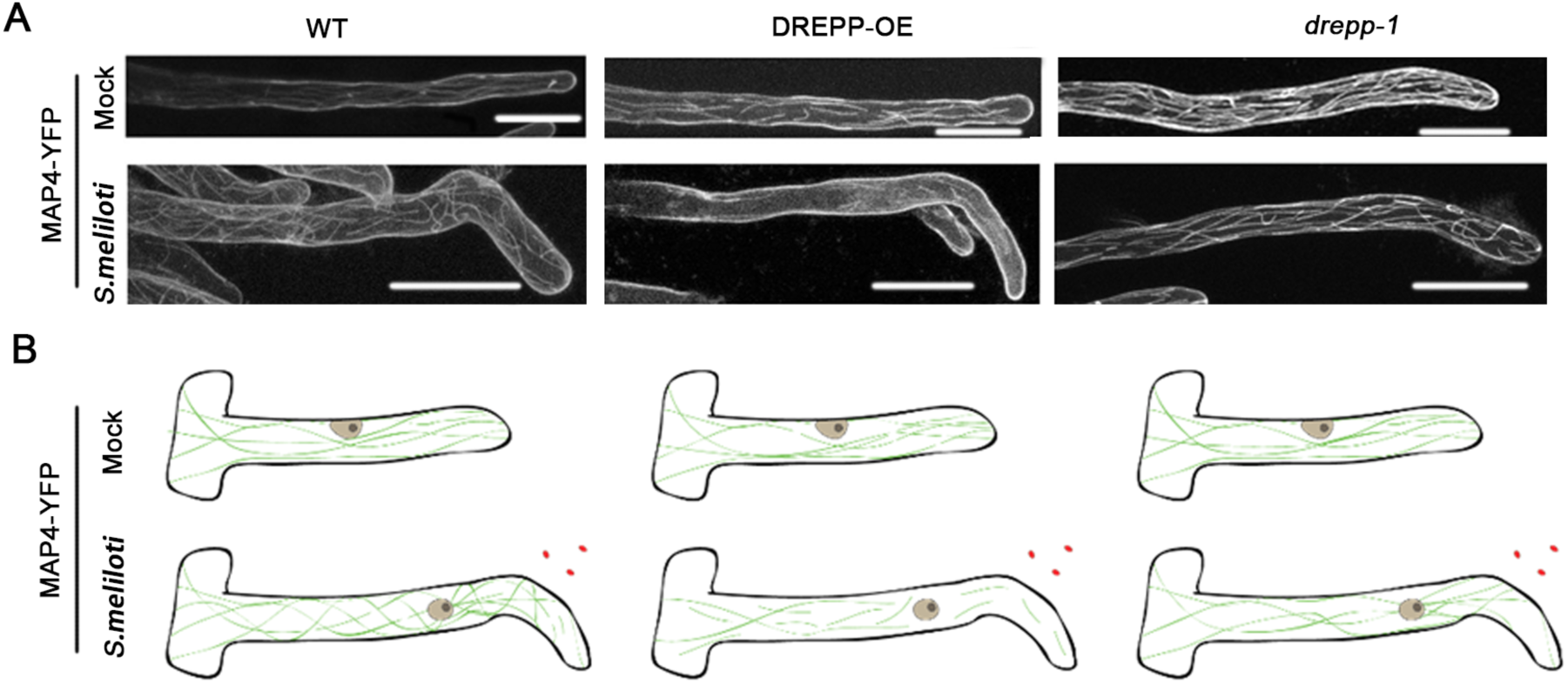
Recruitment of DREPP to nanodomains coincides with microtubule fragmentation. (**A**) *M. truncatula* root hairs of wild-type (A17) were transformed with *pUbi::MAP4-YFP* (WT, left panel) or *pUBI::DREPPGFP//pUBI::MAP4-YFP* (DREPP-OE; middle panel) and *drepp-1* mutant transformed with *pUBI::MAP4-YFP* (*drepp-1*, right panel) and inspected under uninoculated (control) conditions or 7 dpi with *S. meliloti*. Images are maximum intensity z-projections of ∼70 optical sections acquired at 0.5-1 µm increments. Scale bars indicate 20 µm. (**B**) Graphical model of observed microtubule (green) patterns in trichoblasts. Rhizobia and nuclei are indicated in red and grey, respectively.

Since DREPP does not uniformly label MTs we asked whether it associates predominantly with the plus or minus end of the MT filaments and may interfere with the corresponding protein machinery there. For this, we used *N. benthamiana* and co-expressed DREPP together with the plus end markers MICROTUBULE-END-BINDING 1b (EB1b), CLIP-ASSOCIATED PROTEIN (CLASP), the minus end markers GCP3-INTERACTING PROTEIN 1A (GIP1a)/GAMMA-TUBULIN COMPLEX PROTEIN 2 (GCP2)), KATANIN 1 (KTN1), a marker for MT branch points or with SPIRAL 2 (SPR2), which labels both ends and branch points. When performing co-immunoprecipitation experiments using anti-GFP nanobody traps on protein extracts from *N. benthamiana* leaves, we only obtained signals for SPR2 and EB1b (Supplemental Figure 12, line 1 and line 6), indicating that DREPP may predominantly associate with the plus end of the MTs and interferes with their elongation.

Taken together these results show that DREPP controls MT reorganization by interfering with their plus end elongation, predominantly upon localizing to membrane nanodomains after inoculation with rhizobia.

## Discussion

Rhizobial infections of legume root hairs require a tight spatio-temporal coordination of molecular and morphological changes in order to provide a suitable environment for the initial bacterial entry and for maintaining a fast-growing infection thread (IT). Earlier work demonstrated that prolonged exposition of root hairs to isolated NFs or application of rhizobia results in a re-organization of membrane-resident signaling proteins such as the entry receptor LYK3 and the molecular scaffolds FLOT4 and SYMREM1 into membrane nanodomains (Haney et al., 2011; Liang et al., 2018). While the precise molecular composition of this symbiosis-induced nanodomain remains elusive, FLOT4 and actin are ultimately required for the subsequent recruitment of SYMREM1 and LYK3, which has been hypothesized to mark the transition between initial signaling events and morphological changes to prepare the cell for infection (Liang et al., 2018). This process is accompanied by a re-organization of the root hair cytoskeleton (Sieberer et al., 2005). Here, we demonstrate that DREPP is a novel component of this symbiosis-related nanocluster at the PM that is recruited into these domains in an NF-dependent manner (Fig. 2) and by the association with SYMREM1 and FLOT4 (Fig. 4).

In general, root hair growth is characterized by longitudinal cortical MT (CMTs) arrays within the shanks of the RH while the most apical region is devoid of CMTs (Sieberer et al., 2002) in *M. truncatula* but is comprised of rather organelle-depleted cytoplasm with highly enriched secretory vesicles (reviewed in Sieberer et al., 2005). Growth arrest of RHs results in decreased but longitudinally arranged CMTs in *M. truncatula* (Sieberer et al., 2002). In addition to CMTs, legumes display dense arrays of endoplasmic MTs (EMTs) that populate the subapical region between the nucleus and the RH tip (Sieberer et al., 2002; Weerasinghe et al., 2003; Sieberer et al., 2005). The fact that abundant EMTs have not been observed in other model plants such as *A. thaliana* indicates differences in MT organization in between plant families. This is also supported by an additional and interesting difference that has been observed when studying the effect of paclitaxel (taxol), which inhibits MT dynamics by stabilizing and bundling MT polymers (Sieberer et al., 2002). In contrast to *A. thaliana*, (Ketelaar et al., 2002) paclitaxel application to *M. truncatula* roots hairs does not inhibit RH tip growth but only reduces the growth rates (Sieberer et al., 2002) raising the hypothesis whether these EMT arrays serve specialized functions during rhizobial infections of legume root hairs. Given the fragmentation pattern of MTs in root hairs over-expressing DREPP (Fig. 5) and its NF-induced localization into membrane nanodomains (Fig. 2F) we tend to speculate that DREPP serves specialized rather than general functions in *M. truncatula*. This would explain the absence of a generic root hair phenotype in *M. truncatula drepp* mutants (Supplemental Figure S3), which is different from phenotypes observed for the paralogs in *A. thaliana* (Kato et al., 2013). Interestingly, DREPP is conserved in legumes that have lost nodulation such as *Cercis canadensis* or *Nissolia schottii* (Griesmann et al., 2018) indicative of additional functions, likely context dependent. Mechanistically, this functional specialization could be achieved by temporally controlled association to the MT end tracking proteins SPR2 and EB1b (Supplemental Figure S12).

Interestingly, the application of calcium resulted in the same nanodomain localization pattern of DREPP (Supplemental Figure S8). A calcium-dependent interplay between MT association and nanodomain localization has already been reported for other proteins like members of the IQ67 DOMAIN (IQD) family such as IQD24 and IQD25 (Burstenbinder et al., 2017). Although the precise function of these proteins remains unclear, it has been hypothesized that they play role in MT patterning (Abel et al., 2013). However, DREPP recruitment into nanodomains does not seem to be exclusively controlled by NF-induced Ca-spiking as nanodomain accumulation of DREPP was not observed in the LYK3 mutant allele *hcl-4*, a mutant with wild-type like Ca-spiking patterns (Fig. 3; Wais et al., 2000). Furthermore, exogenous application of calcium to *dmi2* and *hcl-1/hcl-4* mutants restored DREPP localization to nanodomains whereas this was not observed in *symrem1* and *flot4* mutants (Supplemental Figure S8). These results led us to conclude that calcium-dependent recruitment to nanodomains is an integral response to active symbiotic signaling that is blocked in *hcl-4* and *dmi2* mutants while SYMREM1 and FLOT4 provide rather structural components for the recruitment. In addition, FLOT4- and SYMREM1-positive nanodomains have been shown to form independent of a disrupted MT network, while interfering with the actin cytoskeleton resulted in a loss of these structures (Liang et al., 2018). By contrast, nanodomain recruitment of several other membrane-resident proteins outside of the symbiosis-specific nanodomain such as HIR1 (Lv et al., 2017), group 1 & 6 remorins and other proteins (Jarsch et al., 2014; Szymanski et al., 2015) have be shown to largely depend on intact MT arrays and/or be restricted in their lateral mobility by them. Our finding that over-expression of DREPP results in altered MT orientation patterns (Supplemental Fig. S4) and that MT fragmentation correlates with the FLOT4- and SYMREM1-dependent nanodomain recruitment of DREPP (Fig. 4; Supplemental Fig. S10) allows the additional hypothesis that this catalytic activity of DREPP may be restricted to membrane nanodomains providing a unique physiological environment. As a consequence, DREPP may serve specialized functions in active MT re-arrangement during root hair curling in response to rhizobial inoculation. Whether DREPP is also involved in MT fragmentation during bacterial release, a process that has been recently described (Kitaeva et al., 2016), remains to be studied in detail but altered or delayed release patterns in 2-weeks old nodules (Supplemental Figure S6) indicates such role.

## Materials and Methods

### Plant growth and phenotypical analysis

*M. truncatula* seeds were activated incubating them in pure sulfuric acid for 10-15 minutes followed by six successive washing steps in sterile water. Seeds surface sterilization was conducted using a 12% Na-hypochlorite solution containing 0.1% SDS for 1 minute followed by extensive washing in sterile water. Seeds were then placed on 1% water agar plates and vernalized at 4°C for 3 days in darkness. Germination was allowed for up to 24h at 24°C in darkness before transferring the seedlings for growth on vertical plates containing Fahraeus medium.

For phenotyping experiments seeds were germinated accordingly. After one day of germination, they were placed onto solid Fahraeus medium (Medicago handbook: https://www.noble.org/medicago-handbook) containing 0.5 mM ammonium nitrate. One week later, the seedlings were transferred onto new Fahraeus medium without nitrate containing 0.1 µM AVG. *M. truncatula* roots were kept for four days under nitrogen starvation conditionsprior to rhizobial inoculation. *S. meliloti* (Sm2011) liquid cultures were sedimented (3 min, 800g), washed and resuspended in liquid FP medium to a final OD_600_ = 0.05-0.07. Each root was covered with 1 ml rhizobia suspension, which was removed after 6 minutes. Afterwards plants were grown at 24°C in white light with the roots being kept in darkness. For growing in pots, the seedlings were directly transferred to a mixture of equal volume of quarz sand and vermiculite and watered with Fahraeus liquid medium (without nitrate) twice a week. After one week growth they were inoculated with *S. meliloti* (Sm2011) at a final OD_600_ = 0.005.

### Hairy Root Transformation

*M. truncatula* hairy root transformation was performed as previously described (Boisson-Dernier et al., 2001). Transgenic *Agrobacterium rhizogenes* (ARqua1) strains, carrying the plasmids of choice, were grown in LB liquid medium for one day and subsequently plated on solid LB agar plates with corresponding antibiotics for another two days before transformation. *M. truncatula* seeds were surface sterilized and vernalized as described above. Emerging roots were removed from the seedlings prior to their shoots being dipped into the *A. rhizogenes* culture and placed on solid Fahraeus medium containing 0.5 mM ammonium nitrate. Transformed seedlings were incubated for three days at 22°C in darkness, followed by 4 days at 22°C in white light with roots covered. One week after transformation, seedlings were transferred onto fresh Fahraeus medium and grown for another 7-9 days at 24°C in white light. Afterwards, they were screened for transformation efficiency using a stereomicroscope scoring the presence of a fluorescent nuclear marker. Untransformed roots were removed and seedlings were transferred onto nitrogen-free solid FP medium containing 0.1 µM AVG. After 3-5 days at 24°C they were inoculated with *S. meliloti* for further studies. All seedling preparation steps were performed under sterile conditions.

### Genotyping of *Tnt1* insertion lines and expression analysis

*M. truncatula* (R108 background) *Tnt1* transposon insertion lines were obtained from the Noble Research Institute (OK, USA). Insertions were verified and precisely mapped using primers listed in Table S3. RNA isolation, cDNA synthesis and quantitative RealTime (RT) PCR were conducted as described earlier (Liang et al., 2018) using a SYBR Green assay (Applied Biosystems). All data were normalized to Ct values of the housekeeping gene ubiquitin (Satge et al., 2016).

### Promoter isolation and analysis

For promoter studies, a 1.5kb long fragment of the putative *DREPP* promoter was amplified from Medicago A17 genome and used to drive expression of *β*-glucuronidase (GUS) in transgenic *M. truncatula* roots. Roots were harvested at 7 days post inoculation with rhizobia and incubated at 37°C in GUS solution (10 mM NaEDTA, 1 mM Potassium ferricyanide, 1 mM Potassium ferrocyanide, 100 mM Sodium phosphate, (pH 7.0), 0,1% Triton-x100 and 0.3% x-Gluc) in darkness. Afterwards they were washed once with ddH2O and stored in 70% ethanol. Analysis of GUS expression was done using stereomicroscopy.

### Construct Design

For expression and purification of the recombinant *M. truncatula* DREPP protein, the coding sequence (GenBank no. XM_013607856.1) was recombined into the Gateway-compatible pDEST17 vector using LR clonase. All other constructs were cloned in a Golden Gate compatible cloning system as previously described (Binder et al., 2014). The DREPP coding and genomic sequences were amplified from mRNA and genomic DNA from A17 wild-type plants, respectively. All the microtubules markers used in Co-IP assays were synthesized by Thermo Scientific. BpiI and BsaI restriction sites were removed from all sequences prior to cloning them into Level II or Level III expression vectors. All designed constructs and cloning primers are listed in Tables S2 and S3, respectively. Identifiers for genes used in this study were list in Table S4.

### DREPP protein expression and purification

The His-tagged DREPP fusion protein was purified from *Escherichia coli* strain BL21 (DE3). Bacteria were grown overnight and then refreshed in 100 ml LB medium containing the respective antibiotics at 37°C. Gene expression was induced at an OD_600_ of 0.6 by adding IPTG (final concentration is 1 mM) in the culture and further grown at 16 °C overnight. Recombinant DREPP was purified using Ni-NTA beads (Qiagen) as described previously (Tao et al., 2010).

### Transformation of *Nicotiana benthamiana* leaves and microsomal fractionation

Transgenic *Agrobacterium tumefaciens* (Agl1), carrying the plasmids of interest, were grown in LB liquid culture overnight at 28°C, using the appropriate antibiotics. The culture was sedimented at 1000g for 2 min and subsequently resuspended in Agromix (10 mM MgCl_2_; 10 mM MES/KOH, pH 5.6; 150 µM Acetosyringone) to a final OD_600_ = 0.3. Bacteria were incubated for 2 hours at 25°C in darkness prior to syringe infiltration into the lower site of *N. benthamiana* leaves. Microsomal fractions protocol was followed as in (Abas and Luschnig, 2010) with some modifications. Extraction buffer was added to frozen root pulverized samples (generally 1.0–1.5 µl EB/mg material, minimum volume 300 µl). The homogenate was transferred to prepared PVPP pellets to adsorb phenolic compounds, vortex, and left for 5 min. Samples were centrifuged twice (600g, 3 min, 4°C) and the pellet (“debris” and nuclear fraction) was kept aside. Supernatants were centrifuge (2,000g, 2 min, 4°C) to obtain a cleared homogenate and the pellet containing organelles was kept aside. Cleared homogenates were diluted with water to 0.37–0.40 M sucrose (usually an equal volume of water was added), vortex and centrifuged (21,000g, 2h, 4 °C) to obtain the microsomal fraction (pellet) and cytosolic fraction (supernatant). For Co-IP assays, DREPP-GFP was co-expressed with the different mCherry-tagged MTs markers in *N. benthamiana* leaves for 2 to 3 days. Extracted proteins were immuno-precipitated using anti-GFP nanobody traps (Chromotek).

### Treatment of *M. truncatula* roots for microscopic analysis

10nM *S. meliloti* Nod Factor and 1mM CaCl_2_ were solubilized in sterile water and applied to excised roots for 12 hours and 2 hours, respectively, prior to their microscopic analysis. For microtubule depolymerization experiments excised roots were emerged in a 10 µM oryzalin solution for 14 hours at room temperature. In all cases the control (mock) samples were treated in the same way using sterile water for incubation. For nodule sections, the material was harvested 2 or 4 weeks after being inoculated with *S. meliloti* in open pots. Nodules were directly embedded in 6 % low melting agarose before being sectioned into semi-thin (70µm) longitudinal sections using a vibratome.

### Whole mount staining of microtubule cytoskeleton

The microtubule cytoskeleton was stained according to Ditengou et al., 2003 with minor modifications. Roots were fixed in 2% paraformaldehyde and 0.25% glutaraldehyde in microtubule-stabilizing buffer (MTSB: 50 mM Pipes, 5 mM EGTA, 5 mM MgSO4, 0,01% Triton X-100, pH 7.0) for 30 min at room temperature (RT). Fixed roots were washed twice 5 min in MTSB and 5 min water, respectively. After partial digestion of cell walls with 0.2% macerozyme and 0.2% driselase in 10 mM MES (pH 5.0) for 30 min at 37°C, roots were incubated in permeablization solution (10 % DMSO/3% NP40 in MTSB), at RT for 1 hr and washed 5 times for 5 min in MTSB at RT. Then, roots were incubated with a monoclonal anti-α-tubulin (bovine), mouse IgG1 antibody (A11126; Molecular Probes), diluted 1:50 in MTSB and supplemented with 2% BSA, overnight at 4 °C. After 2 washes in MTSB (10 min each time at RT), the primary antibody was recognized by an anti-mouse AlexaFluor488 antibody (A-11001; ThermoFisher Scientific) at the final concentration of 1:1000 in MTSB with 2% BSA at RT, in the dark. Roots were successively washed 5 times in MTSB and water (10 min each time) and mounted in ProLong^™^ Gold Antifade solution containing DAPI.

### Confocal Laser-Scanning Microscopy and data analysis

Confocal laser scanning microscopy was performed using a Leica TCS SP8 confocal microscope. Images of cells expressing single fluorophores were acquired using an HC PL APO 20x/0.75 IMM CORR CS2 water immersion objective and the following settings: GFP (Ex: 488 nm; Em: 500-550 nm); YFP (Ex: 514 nm; Em: 520-552 nm); mCherry (Ex: 561 nm; Em: 575-630 nm). Samples co-expressing two fluorophores were imaged in sequential mode between frames. Due to the low signal intensities of the MAP4-YFP reporter, the corresponding fluorescence was detected by using Leica HyD detectors. Images were taken with a Leica DFC350FX digital camera. Microtubule orientation analysis was done using CytoSpectre (Kartasalo et al., 2015) and all other images analyses and projections were performed with ImageJ/(Fiji) (Schindelin et al., 2012). For kymographs analysis, images were acquired over 5 min. Regions of interest were manually selected and analyzed by using the kymograph plugin in Fiji. For nanodomain perimeter analysis, images were subjected to background substraction with a rolling ball radius of 20 pixels prior to analysis.

### Lipid binding assays

Lipid binding assays were performed as suggested by the supplier. In brief, PIP Strips™ (Echelon Biosciences, Salt Lake City, UT, USA) were blocked using PBST (10 mM Tris–HCl (pH 8.0), 150 mM NaCl and 0.1% (w/v) Tween-20) supplemented with 3% fatty-acid free bovine serum albumin (BSA) for 1 hour at 25°C. The strips were then incubated with recombinant DREPP protein in 7 ml PBST (final concentration: 100 ng/ml) at 4°C overnight before being washed with PBST 5 times (5 minutes per wash), blotted and detected using an anti-His antibody (Sigma) for 2 hours at 25°C. The blots were subsequently washed 5 times in PBST prior to visualization.

### Microtubule binding assays

The co-sedimentation of the recombinant DREPP protein and microtubules was performed as previously published method (Li et al., 2011) with the following modifications: purified protein preparations were sedimented at 200,000 *g* and 4°C for 20 min before use. Pre-polymerized and paclitaxel stabilized microtubules were obtained using brain tubulin (2mg/ml; stored at −80°C), which was incubated in PEM buffer containing paclitaxel (PEMT buffer; 1 mM MgCl_2_, 1 mM EGTA, 100 mM PIPES-KOH, 1mM GTP, 150mM KCl and 10 mM paclitaxel, pH 6.9) at 30°C for 20 minutes. Different amounts of recombinant DREPP fusion protein were added and the mixtures were incubated at 30°C for an additional 20 minutes. After sedimentation at 100,000 *g* for 20 minutes, supernatant and pellets were subjected to SDS-PAGE and DREPP was detected using anti-His antibodies (Sigma).

### Phylogenetic analysis

DREPP orthologs were searched within 104 plant genomes covering all main orders from Gymnosperms to Angiosperms (See Table S1). Searches were performed using tBLASTn v2.8.1+ (Camacho et al., 2009), an e-value of 1e-10 and the DREPP protein from *Medicago truncatula* as query. From blast results, protein sequences of all putative orthologs were retrieved and aligned using MUSCLE v3.8.31 (Edgar, 2004) with default parameters. Then, the alignment was cleaned by removing all positions containing more than 20% of gaps using trimAl v1.5rev57 (Capella-Gutierrez et al., 2009). Resulting alignments were subjected to phylogenetic analysis using Maximum Likelihood approach and the IQ-TREE v1.6.7 (Nguyen et al., 2015). Prior to the phylogenetic analysis, the evolutionary best fitting model was tested using ModelFinder (Kalyaanamoorthy et al., 2017) and retained according to the Bayesian Information Criteria. Branch support was tested with 10,000 replicates of UltrFast Bootstraps (Hoang et al., 2018). The final tree was visualized and annotated using the iTOL platform v4.3 (Letunic and Bork, 2016).

DREPP functional domains (PF05558.12) were retrieved for each ortholog using the Pfam database (Finn et al., 2016), an evalue of 0.01 and default parameters. Domains were then mapped on the phylogenetic tree using iTOL.

## Supplemental Data

**Supplemental Figure S1.** Members of the DREPP protein family can be found throughout the angiosperms.

**Supplemental Figure S2.** Transcriptional regulation of DREPP.

**Supplemental Figure S3.** Medicago *drepp* mutants do not display a generic nucleus positioning phenotype in root hairs.

**Supplemental Figure S4.** Cortical microtubules patterns in root epidermal cells of different genotypes.

**Supplemental Figure S5.** DREPP over-expression (DREPP-OE) reduces rhizobial infections.

**Supplemental Figure S6.** Patterning within Medicago *drepp* mutant nodules is altered.

**Supplemental Figure S7.** A bifunctional mechanism of PM association for DREPP.

**Supplemental Figure S8.** Calcium treatment induces DREPP-GFP re-localization into membrane nanodomains.

**Supplemental Figure S9.** DREPP directly associates with microtubules.

**Supplemental Figure S10.** DREPP nanodomains are linked with microtubule fragmentation in root hairs.

**Supplemental Figure S11.** Depolymerized microtubule filaments is not affecting DREPP-GFP location to PM.

**Supplemental Figure S12.** Using Co-Immunoprecipitation (Co-IP) searching for DREPP interact proteins.

**Supplemental Table S1.** Genome information that was used to generate the phylogenetic tree of DREPP.

**Supplemental Table S2.** Constructs used in this study.

**Supplemental Table S3.** List of primers used for this study.

**Supplemental Table S4.** Identifiers for genes used in this study.

## Acknowledgements

This study was supported by the German Research Foundation DFG) under Germany’s Excellence Strategy (CIBSS – EXC-2189 – Project ID 39093984), through a sub-grant to the University of Freiburg (T.O.) made by the University of Cambridge through the Engineering Nitrogen Symbiosis for Africa (ENSA) project, which is funded by the Bill & Melinda Gates Foundation (OPP1172165) and the China Scholarship Council (CSC) (grant no. 201708080016 to CS). JK and PMD belong to the LRSV, which is part of the TULIP LABEX (ANR-10-LABX-41). We thank the staff of the Life Imaging Center (LIC) in the Center for Biological Systems Analysis (ZBSA) of the Albert-Ludwigs-University Freiburg for help with their confocal microscopy resources, and the excellent support in image recording. Special thanks also to Giles Oldroyd (SLCU Cambridge, UK) for providing the *S. meliloti* Nod factors and to Magda Magiera (Carsten Janke group, Institut Curie, PSL Research University, France) for providing tubulin for the co-sedimentation assay. The *Medicago truncatula* plants utilized in this research project, which are jointly owned by the Centre National De La Recherche Scientifique, were obtained from Noble Research Institute, LLC and were created through research funded, in part, by a grant from the National Science Foundation, NSF-0703285.

Furthermore, we would like to thank Eija Schulze for her excellent technical support and Beatrice Lace illustrating the model. Also, many thanks all the members of our team for fruitful discussions and providing their individual expertise throughout the course of the project.

## Author contributions

C.S., M.L.K., T.O. designed the research; C.S., M.L.K., C.H-R., M.B. F.A.D. J.K. performed research; J.K., P.M.D. contributed new computational tools; C.S., P.M.D., analyzed data; C.S., T.O. wrote the paper

