## Supplemental Material for "The *Medicago truncatula* DREPP protein triggers microtubule fragmentation in membrane nanodomains during symbiotic infections"

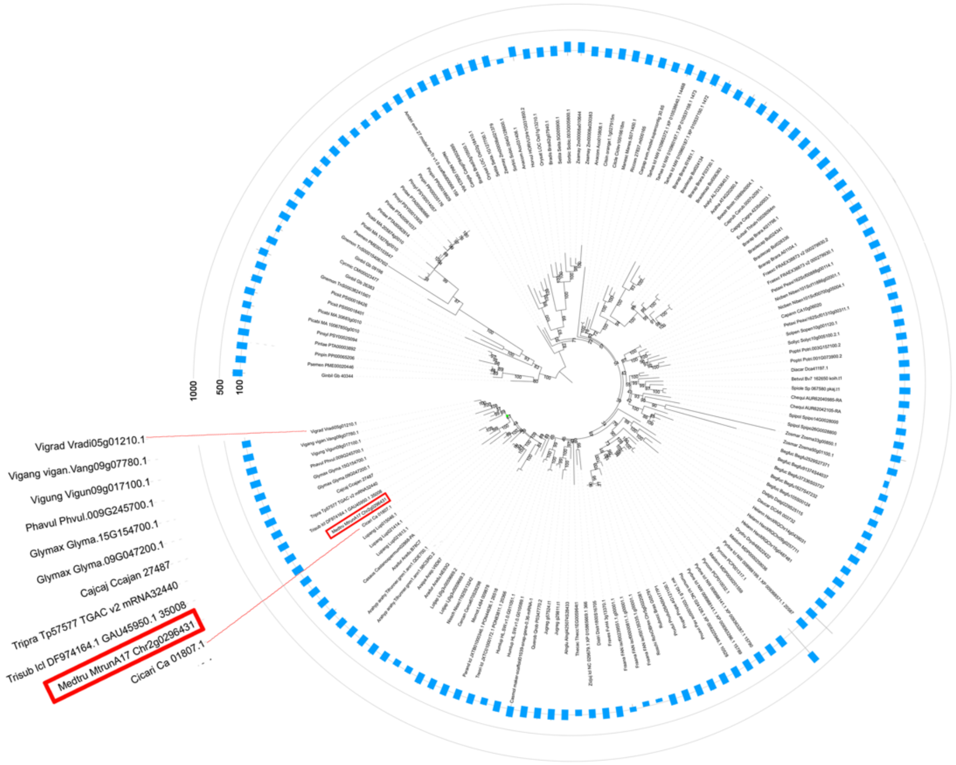


**Supplemental Figure S1. Members of the DREPP protein family can be found throughout the angiosperms.** Maximum likelihood tree of DREPP orthologs (model: Dayhoff+F+R5). The tree was rooted on Gymnosperms. UltraFast Bootstraps are indicated for each node. DREPP functional PFAM domains (blue, PF05558.12) are mapped in the outer circle and represented by a blue rectangle. Length of the rectangle is proportional to the length of the domain. Red box indicates the Medicago *DREPP* gene. The annotation for DREPP (Chr2g0296431) in this tree is based on MtrunA17r5.0-ANR (https://medicago.toulouse.inra.fr/MtrunA17r5.0-ANR/), which corresponds to Medtr2g437530 used in Phytozome 12 (https://phytozome.jgi.doe.gov/pz/portal.html) or the *M. truncatula* Genome Database (http://www.medicagogenome.org/).

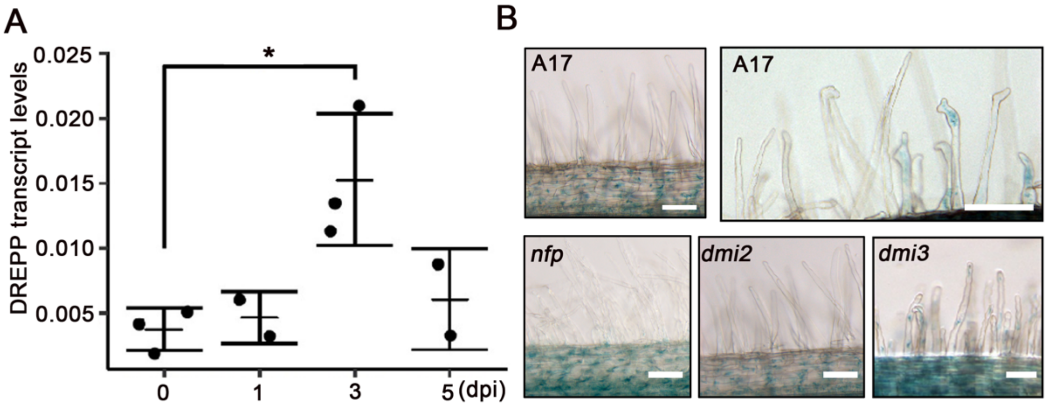


**Supplemental Figure S2. Transcriptional regulation of *DREPP*.** (**A**) Endogenous *DREPP* transcripts were assessed by qRT-PCR at different timepoints after inoculation of *M. truncatula* roots with *S. meliloti*. Transcript levels were normalized to Ubiquitin. Data are given as mean ± SE. Statistics were performed by using an unpaired, two-tailed Student t-test (and nonparametric tests). *= $p<0.05$. (**B**) The putative *DREPP* promoter (1.5 kb upstream of the translational start) was tested using a *pDREPP::GUS* reporter construct in wild-type (A17) and different symbiosis mutants backgrounds. GUS staining were performed at 7dpi with *S. meliloti*. Scale bar indicates 50 µm.

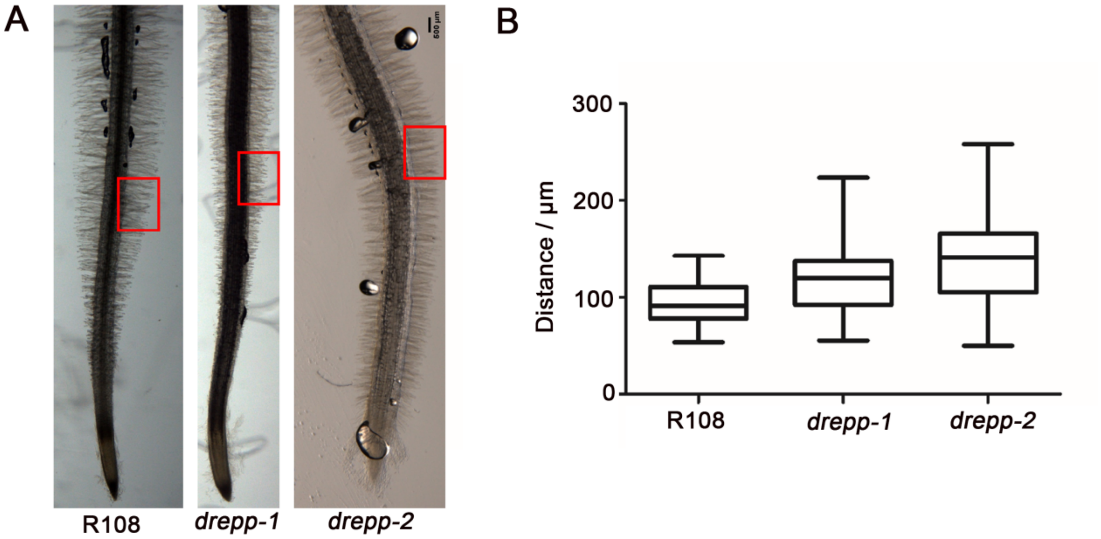


**Supplemental Figure S3. Medicago *drepp* mutants do not display a generic nucleus positioning phenotype in root hairs.** (**A**) Images were taken from seedlings grown on Fahraeus medium for 7 days. Representative positions of scored root hairs about 10mm above the root tip are marked by a red box. Scale bars indicates 500µm. (**B**) Distance between the nucleus and root hair tip after DAPI staining. n= 4 independent roots. Data are given as mean ± SE. Statistics were performed using an unpaired, two tailed Student t-test (and nonparametric tests). No significant differences were observed.

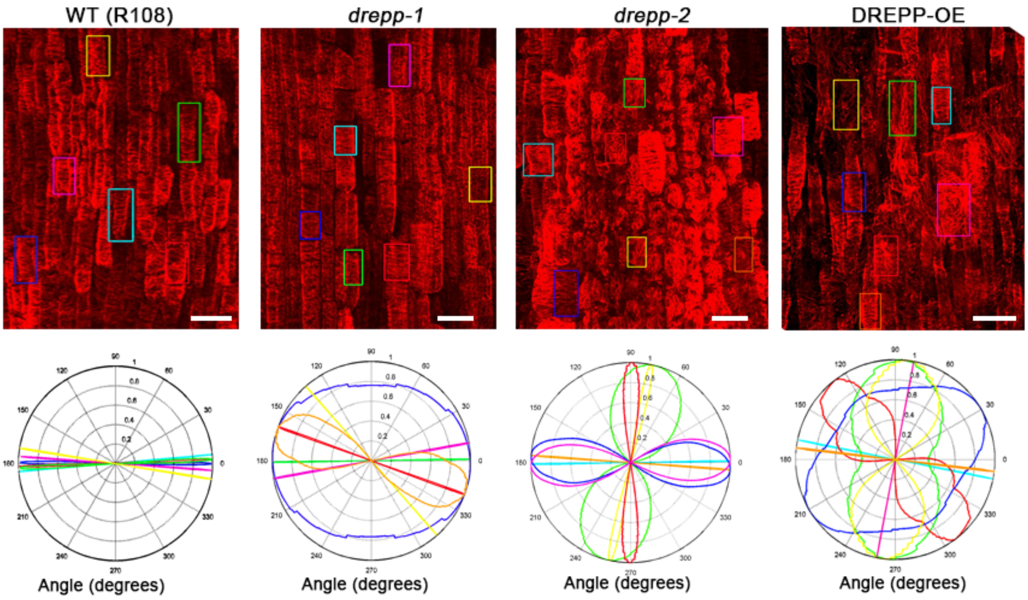


**Supplemental Figure S4. Cortical microtubules patterns in root epidermal cells of different genotypes.** For R108, *drepp-1* and *drepp-2* mutant and DREPP-overexpressing (DREPP-OE) plants, seedlings were grown on Fahraeus medium for 7 days (or 5 days (DREPP-OE)) prior to immunofluorescence experiments using anti-tubulin antibodies. Scale bars indicate 15µm. The boxes indicate the cells analyzed for microtubule orientation using the CytoSpectre software (Kartasalo et al., 2015). Comparable results were obtained for other images.

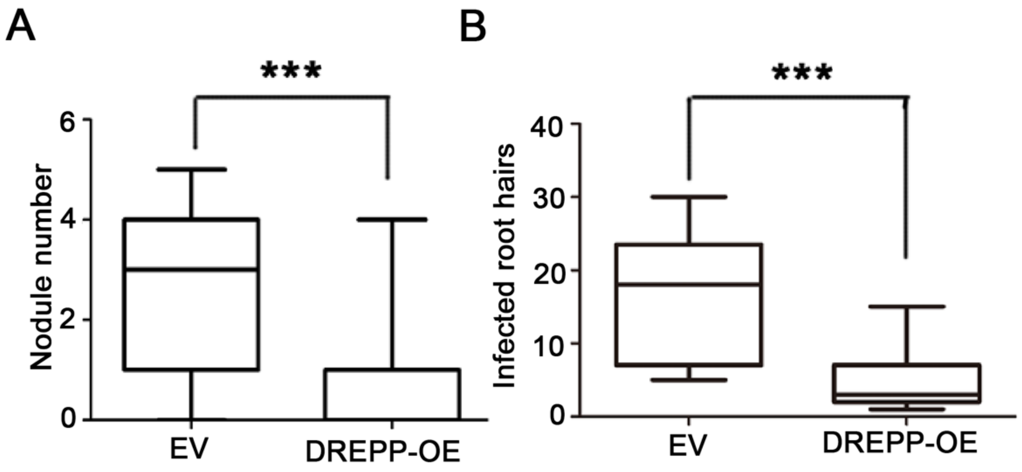


**Supplemental Figure S5. DREPP over-expression (DREPP-OE) reduces rhizobial infections**. Nodules (**A**) and number of infected root hairs (curled root hairs) (**B**) were scored at 7 dpi with n >=15 root systems for each genotype and condition. Data are given as mean ± SE. Statistics were performed by using an unpaired, two-tailed Student t-test (and nonparametric tests). ***= $p < 0.001$. EV= empty vector control.

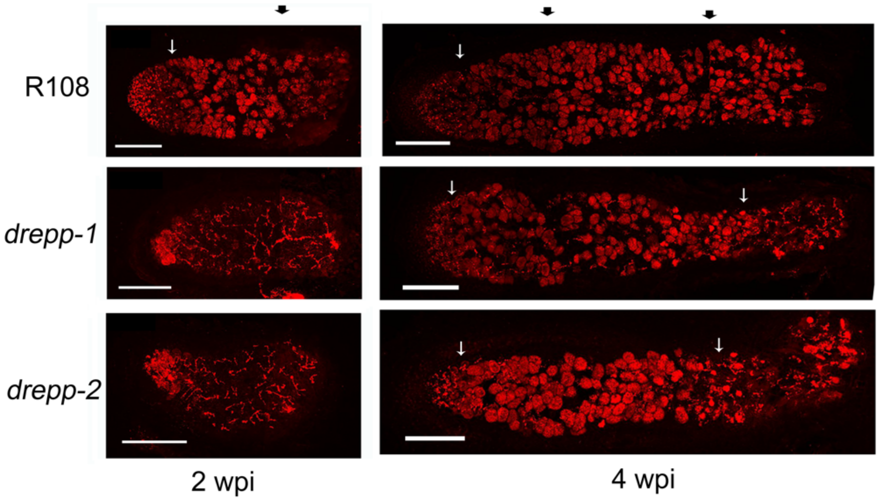


**Supplemental Figure S6. Patterning within Medicago *drepp* mutant nodules is altered.** Semi-thin (70µm) longitudinal sections of 2- and 4-weeks old whole nodules from R108, *drepp-1* and *drepp-2* plants grown in open pots. Red color indicates mCherry-labeled rhizobia. Scale bars indicate 150µm. Black arrows indicate the images stitching region for panorama. The white arrows indicate the transition between zone II to III (left) and III to IV (right).

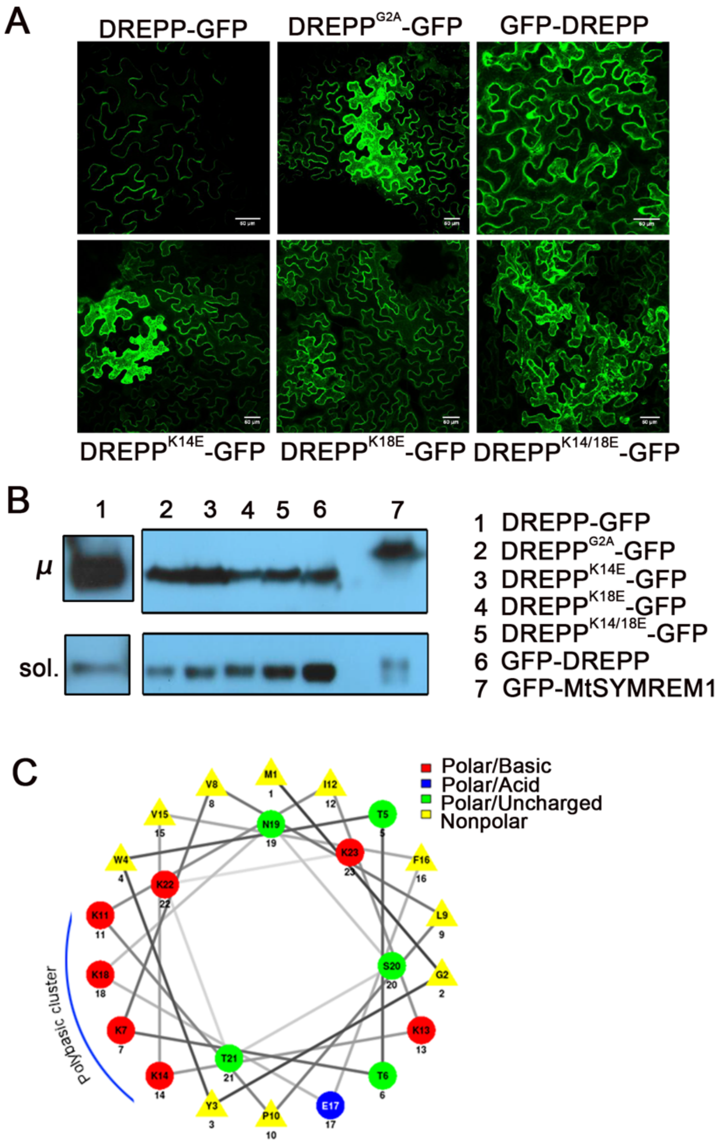


**Supplemental Figure S7. A bifunctional mechanism of PM association for DREPP.** (**A**) Different DREPP variants were expressed in *N. benthamiana* leaf epidermal cells. C-terminal fusion of GFP (DREPP-GFP) resulted in PM association while this was decreased in the myristoylation mutant (G2A) or by fusing GFP to the N-terminus of DREPP (GFP-DREPP). Moreover, replacing core lysine (K) residues by glutamate (E) within the predicted polybasic cluster (**C**) resulted in increased cytosolic fluorescence (**A**, lower row). Scale bars indicate 50 µm. (**B**) Microsomal fractions (µ) of samples shown above indicate increased soluble (sol.) fractions in the mutant variants. PM-localized *M. truncatula* SYMREM1 (MtSYMREM1) was used as a control. Due to the comparably low expression of non-mutated DREPP-GFP, the corresponding Western Blots required increased exposition times. (**C**) Prediction of a polybasic cluster within an N-terminal 23 amino acid long stretch of the DREPP protein using the NetWheels server (http://lbqp.unb.br/NetWheels). F= phenylalanine, N= asparagine, M= methionine, P= proline, S= serine, T= threonine, Y= tyrosine. Numbers indicate positions in the DREPP protein.

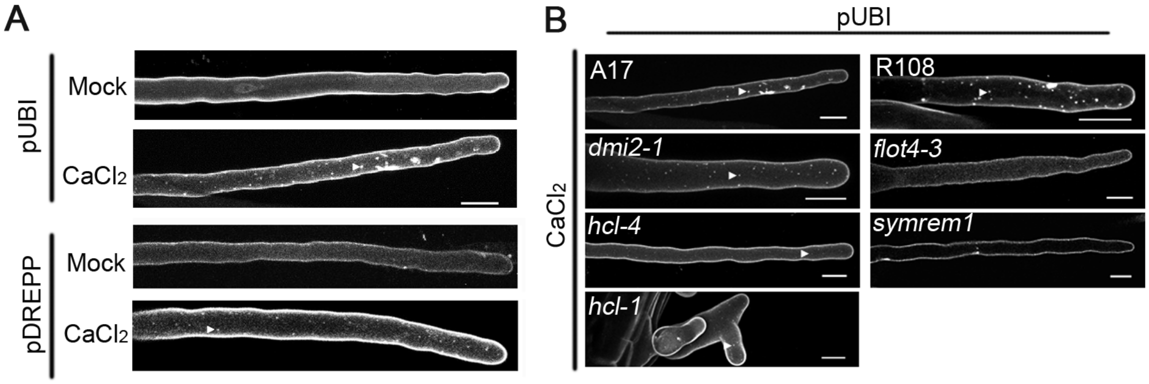


**Supplemental Figure S8. Calcium treatment induces DREPP-GFP relocalization into membrane nanodomains.** *M. truncatula* roots of indicated mutant seedlings and wild-type (R108, A17) seedlings expressing DREPP under the control of the Ubiquitin (pUBI::DREPP-GFP) or the native DREPP promoter (pDREPP::DREPP-GFP). Confocal microscopy was used to image fluorescence of GFP fusion proteins, z-stacks are shown as maximum intensity projections. Root hairs were investigated after calcium treatment (2h, 1mM $CaCl_2$). Nanodomains are indicated by arrowheads. Scale bars = 20 µm.

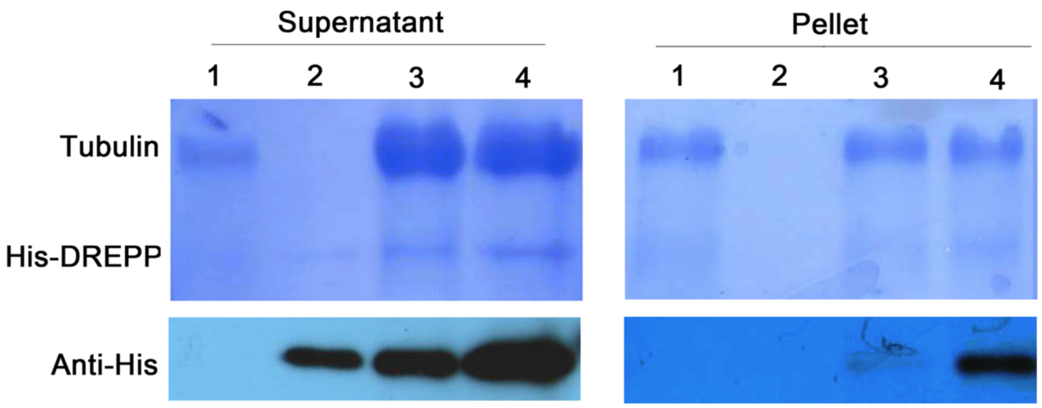


**Supplemental Figure S9. DREPP directly associates with microtubules.** An *in vitro* co-sedimentation assay was used to determine direct DREPP binding to microtubules. Coomassie Blue stained SDS-PAGE (upper panels) indicate presence of the recombinant proteins. His-DREPP was detected by Western Blot analysis (lower panels) using an anti-His antibody. The clear band in lane 4 (Western blot) of the pellet fraction indicates direct microtubule association. Lane 1: 20μg tubulin; Lane 2: 1.5 μg His-DREPP; Lane 3: 20μg tubulin + 2.5 μg His-DREPP; Lane 4: 20μg tubulin + 5 μg His-DREPP.

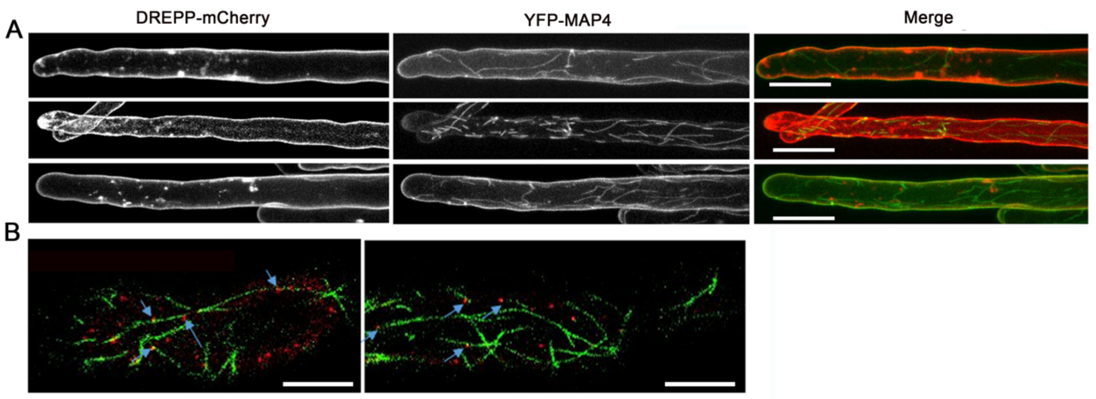


**Supplemental Figure S10. DREPP nanodomains are linked with microtubule fragmentation in root hairs.** (**A**) Three selected *M. truncatula* (A17) root hairs co-expressing DREPP-mCherry and the microtubule marker YFP-MAP4 at 7 dpi with *S. meliloti*. All root hairs showed microtubule fragmentation upon rhizobial inoculation that coincided with nanodomain-localized DREPP (blue arrows) (**B**). Scale bars indicate 20 µm (**A**) and 10 µm (**B**).

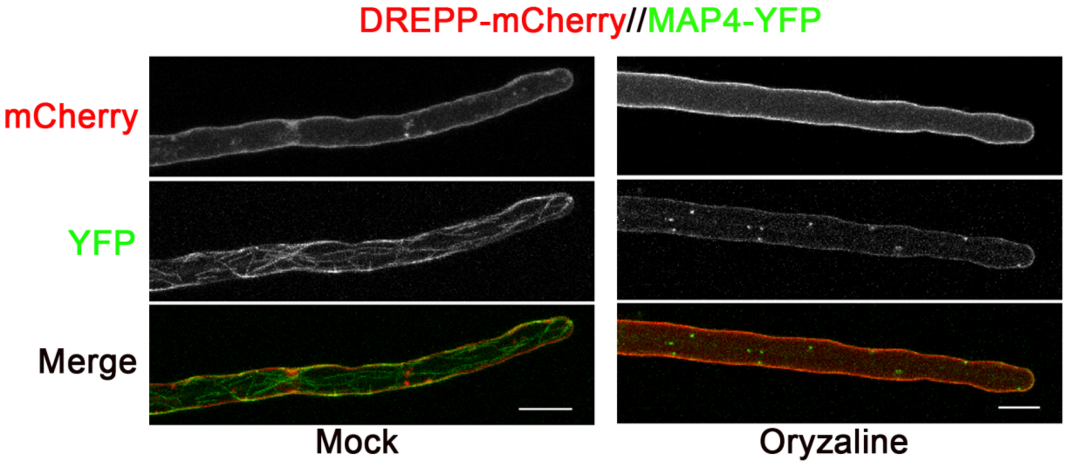


**Supplemental Figure S11. Depolymerization of microtubule filaments does not affect DREPP-GFP localization.** *M. truncatula* roots were transformed with a ProUbi-DREPP-GFP//ProUbi-MAP4-YFP construct. Confocal microscopy was used for image acquisition. Root hairs were investigated (left) under mock ($H_2O$) conditions and (right) upon Oryzaline treatment that induced microtubule depolymerization. Scale bars = 20 µm.

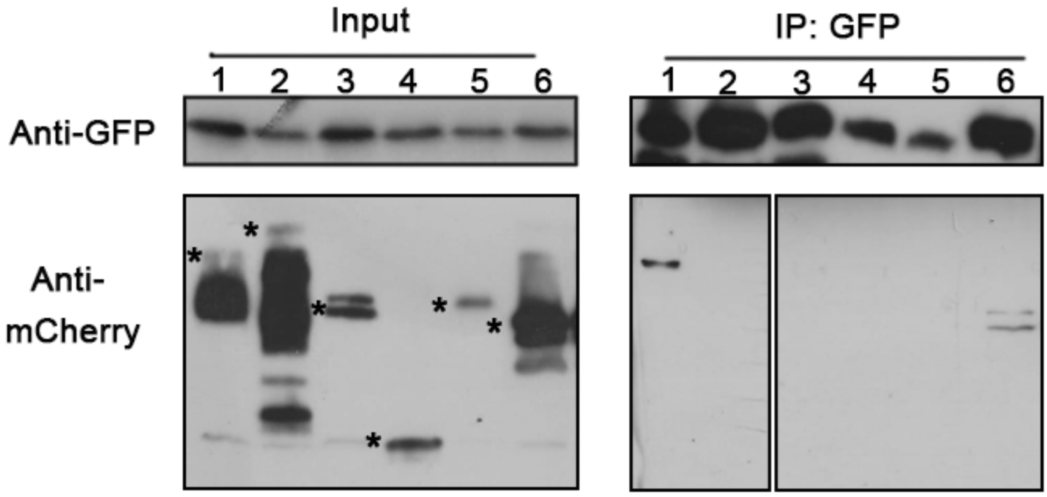


**Supplemental Figure S12. Identifying DREPP-interacting proteins by co-immunoprecipitation (co-IP).** DREPP-GFP was co-expressed with a series of microtubule-associated proteins in *N. benthamiana* leaf epidermal cells. Samples were harvested 3 days after co-infiltration. Co-IP experiments were performed using anti-GFP nanobody traps and immuno-precipitated proteins were detected using an α-mCherry antibody. Lanes: 1-DREPP-GFP + SPR2-mCherry; 2-DREPP-GFP + mCherry CLASP; 3-DREPP-GFP + KTN1-mCherry; 4 DREPP-GFP + GIP1a-mCherry; 5- DREPP-GFP + GCP2-mCherry; 6- DREPP-GFP + EB1a-mCherry.

**Table S1: Genome information that was used to generate the phylogenetic tree of DREPP.**

| Abbreviation | Species | Order | Family | Genome version | Reference |
|---|---|---|---|---|---|
| Alnglut | Alnus glutinosa | Fagales | Betulaceae | 1 | 10.1126/science.aat1743 |
| Ambtri | Amborella trichopoda | Amborellales | Amborellaceae | 1.0 | 10.1126/science.1241089 |
| Anacom | Ananas comosus | Bromeliales | Bromeliaceae | 3.0 | 10.1038/ng.3435 |
| Arahal | Arabidopsis halleri | Brassicales | Brassicaceae | 1.1 | unpublished |
| Aralyr | Arabidopsis lyrata | Brassicales | Brassicaceae | 2.1 | 10.1038/ng.807 |
| Aratha | Arabidopsis thaliana | Brassicales | Brassicaceae | TAIR10 | 10.1093/nar/gkr1090 |
| Aradur | Arachis duranensis | Fabales | Fabaceae | NC | 10.1038/ng.3517 |
| Arahyp | Arachis hypogaea | Fabales | Fabaceae | 1.0 | unpublished |
| Araipa | Arachis ipaensis | Fabales | Fabaceae | NC | 10.1038/ng.3517 |
| Begfuc | Begonia fuchsioides | Cucurbitales | Begoniaceae | 1.0 | 10.1126/science.aat1743 |
| Betvul | Beta vulgaris ssp. vulgaris KWS2320 | Caryophyllales | Amaranthaceae | NC | 10.1038/nature12817 |
| Boestr | Boechera stricta | Brassicales | Brassicaceae | 1.2 | unpublished |
| Bradis | Brachypodium distachyon | Poales | Poaceae | 3.1 | 10.1038/nature08747 |
| Braolecap | Brassica oleraceae capitata | Brassicales | Brassicaceae | 1.0 | 10.1038/ncomms4930 |
| Brarap | Brassica rapa FPsc | Brassicales | Brassicaceae | 1.3 | unpublished |
| Cajcaj | Cajanus cajan | Fabales | Fabaceae | 1.0 | 10.1038/nbt.2022. |
| Capgra | Capsella grandiflora | Brassicales | Brassicaceae | 1.1 | 10.1038/ng.2669 |
| Caprub | Capsella rubella | Brassicales | Brassicaceae | 1.0 | 10.1038/ng.2669 |
| Capann | Capsicum annuum cvCM334 | Solanales | Solanaceae | 1.55 | doi.org/10.1038/ng.2877 |
| Carpap | Carica papaya | Brassicales | Caricaceae | ASGPBV0.4 | 10.1038/nature06856 |
| Casmol | Castanea mollissima | Fagales | Fagaceae | 1.1 | unpublished |
| Casaus | Castanospermum australe | Fabales | Fabaceae | 1.0 | unpublished |
| Casgla | Casuarina glauca | Fagales | Casuarinaceae | 1.0 | 10.1126/science.aat1743 |
| Cercan | Cercis canadensis | Fabales | Caesalpiniaceae | 1.0 | 10.1126/science.aat1743 |
| Chafas | Chamaecrista fasciculata | Fabales | Fabaceae | 1.0 | 10.1126/science.aat1743 |
| Chequi | Chenopodium quinoa | Caryophyllales | Amaranthaceae | 1.0 | 10.1038/nature21370 |
| Cicari | Cicer arietinum ICC4958 | Fabales | Fabaceae | 1.1 | 10.1038/srep12806 |
| Citlan | Citrullus lanatus subsp. vulgaris 97103 | Cucurbitales | Cucurbitaceae | 1.0 | 10.1038/ng.2470 |
| Citcle | Citrus clementina | Sapindales | Rutaceae | 1.0 | 10.1038/nbt.2906 |

| | | | | | |
|---|---|---|---|---|---|
| Citsin | Citrus sinensis | Sapindales | Rutaceae | 1.1 | 10.1038/nbt.2906 |
| Cucmel | Cucumis melo | Cucurbitales | Cucurbitaceae | 4.0 | doi.org/10.1073/pnas.1205415109 |
| Cucsat | Cucumis sativus PI183967 | Cucurbitales | Cucurbitaceae | 1.0 | 10.1038/ng.2801 |
| Cucmax | Cucurbita maxima | Cucurbitales | Cucurbitaceae | 1.1 | https://doi.org/10.1016/j.molp.2017.09.003 |
| Cucmos | Cucurbita moschata | Cucurbitales | Cucurbitaceae | 1.0 | https://doi.org/10.1016/j.molp.2017.09.004 |
| Cucpep | Cucurbita pepo | Cucurbitales | Cucurbitaceae | 4.1 | unpublished |
| Cycmic | Cycas micholitzii | Cycadales | Cycadaceae | 1.0 | unpublished |
| Datglo | Datisca glomerata | Cucurbitales | Dasticaceae | 1.0 | 10.1126/science.aat1743 |
| Daucar | Daucus carota | Apiales | Apiaceae | 2.0 | 10.1038/ng.3565 |
| Diacar | Dianthus caryophyllus | Caryophyllales | Caryophyllaceae | 1.0 | doi:10.1093/dnares/dst053 |
| Distri | Discaria trinervis | Rosales | Rhamnaceae | 1.0 | 10.1126/science.aat1743 |
| Drydru | Dryas drummondii | Rosales | Rosaceae | 1.0 | 10.1126/science.aat1743 |
| Eutsal | Eutrema salsugineum | Brassicales | Brassicaceae | 1.0 | 10.3389/fpls.2013.00046 |
| Fraves | Fragaria vesca | Rosales | Rosaceae | 4.0 | 10.1093/gigascience/gix124 |
| Fraana | Fragaria x ananassa | Rosales | Rosaceae | 1.1 | unpublished |
| Fraxexc | Fraxinus excelsior | Lamiales | Oleaceae | 2.0 | doi:10.1038/nature20786 |
| Ginbil | Ginkgo biloba | Ginkgoales | Ginkgoaceae | 1.0 | 10.1186/s13742-016-0154-1 |
| Glymax | Glycine max | Fabales | Fabaceae | Wm82.a2.v1 | 10.1038/nature08670 |
| Gnemon | Gnetum montanum | Gnetales | Gnetaceae | 1.0 | 10.5061/dryad.0vm37 |
| Gosrai | Gossypium raimondii | Malvales | Malvaceae | 2.1 | 10.1038/nature11798 |
| Helann | Helianthus annuus | Asterales | Asteraceae | r1.2 | 10.1038/nature22380 |
| Horvul | Hordeum vulgare | Poales | Poaceae | 1.0 | 10.1038/nature22043 / 10.1038/sdata.2017.44 |
| Humlup | Humulus lupulus | Rosales | Cannabaceae | 1.1 | 10.1093/pcp/pcu169 |
| Jugreg | Juglans regia | Fagales | Juglandaceae | 1.1 | 10.1111/tpj.13207 |
| Lagsic | Lagenaria siceraria | Cucurbitales | Cucurbitaceae | 1.0 | 10.1111/tpj.13722 |
| Lotjap | Lotus japonicus | Fabales | Fabaceae | 3.0 | 10.1093/dnares/dsn008 |
| Lupang | Lupinus angustifolius | Fabales | Fabaceae | 1.0 | 10.1111/pbi.12615 |
| Maldom | Malus domestica | Rosales | Rosaceae | 3.0 | unpublished |
| Manesc | Manihot esculenta | Malpighiales | Euphorbiaceae | 7.0 | 10.1038/nbt.3535 |
| Medtru | Medicago truncatula | Fabales | Fabaceae | 5.0 | 10.1038/s41477-018-0286-7 |
| Mimpud | Mimosa pudica | Fabales | Fabaceae | 1.0 | 10.1126/science.aat1743 |
| Momcha | Momordica charantia | Cucurbitales | Cucurbitaceae | NC | unpublished |
| Mornot | Morus notabilis | Rosales | Moraceae | 2.0 | 10.1038/ncomms3445 |
| Nelnuc | Nelumbo nucifera | Proteales | Nelumbonaceae | 1.0 | 10.1186/gb-2013-14-5-r41 |

| | | | | | |
|---|---|---|---|---|---|
| Nicben | Nicotiana benthamiana | Solanales | Solanaceae | 1.0.1 | doi.org/10.1094/MPMI-06-12-0148-TA |
| Nissch | Nissolia schottii | Fabales | Fabaceae | 1.0 | 10.1126/science.aat1743 |
| Orysat | Oryza sativa | Poales | Poaceae | 7.0 | 10.1093/nar/gkl976 |
| Parand | Parasponia andersonii | Rosales | Cannabaceae | 1.0 | 10.1073/pnas.1721395115 |
| Petaxi | Petunia axillaris | Solanales | Solanaceae | 1.6.2 | 10.1038/nplants.2016.74 |
| Phavul | Phaseolus vulgaris | Fabales | Fabaceae | 2.0 | 10.1038/ng.3008 |
| Picabi | Picea abies | Pinales | Pinaceae | 1.0 | 10.1038/nature12211 |
| Picglau | Picea glauca | Pinales | Pinaceae | 4.1 | doi.org/10.1093/bioinformatics/btt178 |
| Picsit | Picea sitchensis | Pinales | Pinaceae | 1.0 | unpublished |
| Pinpin | Pinus pinaster | Pinales | Pinaceae | 1.0 | unpublished |
| Pinsyl | Pinus sylvestris | Pinales | Pinaceae | 1.0 | unpublished |
| Pintae | Pinus taeda | Pinales | Pinaceae | 1.0 | 10.1534/genetics.113.159715 |
| Poptri | Populus trichocarpa | Malpighiales | Salicaceae | 3.1 | 10.1126/science.1128691 |
| Pruavi | Prunus avium | Rosales | Rosaceae | 1.0 | 10.1093/dnares/dsx020 |
| Prudul | Prunus dulcis | Rosales | Rosaceae | 2.0 | unpublished |
| Prumum | Prunus mume | Rosales | Rosaceae | 1.0 | unpublished |
| Pruper | Prunus persica | Rosales | Rosaceae | 2.1 | 10.1038/ng.2586 |
| Psemen | Pseudotsuga menziesii | Pinales | Pinaceae | 1.0 | 10.1534/g3.117.300078 |
| Pyrcom | Pyrus communis | Rosales | Rosaceae | 1.1 | 10.1371/journal.pone.0092644 |
| Pyrbre | Pyrus x bretschneideri | Rosales | Rosaceae | 1.0 | unpublished |
| Querob | Quercus robur | Fagales | Fagaceae | 1.0 | 10.1111/1755-0998.12425 |
| Riccom | Ricinus communis | Malpighiales | Euphorbiaceae | 0.1 | 10.1038/nbt.1674 |
| Roschi | Rosa chinensis | Rosales | Rosaceae | 2.0 | 10.1038/s41588-018-0110-3 |
| Rubocc | Rubus occidentalis | Rosales | Rosaceae | 1.0 | doi.org/10.1111/tpj.13215 |
| Setita | Setaria italica | Poales | Poaceae | 2.2 | 10.1038/nbt.2196 |
| Sollyc | Solanum lycopersicum | Solanales | Solanaceae | ITAG2.4 | 10.1038/nature11119 |
| Solpen | Solanum pennellii | Solanales | Solanaceae | 2.0 | doi.org/10.1038/ng.3046 |
| Sorbic | Sorghum bicolor | Poales | Poaceae | 3.1.1 | 10.1111/tpj.13781 |
| Spiole | Spinach oleracea | Caryophyllales | Amaranthaceae | NC | unpublished |
| Spipol | Spirodela polyrhiza | Alismatales | Araceae | 2.0 | 10.1038/ncomms4311 |
| Tarhas | Tarenaya hassleriana | Brassicales | Cleomaceae | 1.0 | 10.1105/tpc.113.113480 |
| Thecac | Theobroma cacao | Malvales | Malvaceae | 1.1 | 10.1186/gb-2013-14-6-r53 |
| Treori | Trema orientalis | Rosales | Cannabaceae | 1.0 | 10.1073/pnas.1721395115 |
| Tripra | Trifolium pratense | Fabales | Fabaceae | 2.0 | 10.1038/srep17394 |

| | | | | | |
|---|---|---|---|---|---|
| Trisub | Trifolium subterraneum | Fabales | Fabaceae | 1.0 | unpublished |
| Vigang | Vigna angularis | Fabales | Fabaceae | 1.0 | 10.1038/srep080669 |
| Vigrad | Vigna radiata | Fabales | Fabaceae | 1.0 | 10.1038/ncomms6443 |
| Vigung | Vigna unguiculata | Fabales | Fabaceae | 1.1 | unpublished |
| Zeamay | Zea mays PH207 | Poales | Poaceae | 1.1 | 10.1105/tpc.16.00353 |
| Zizjuj | Ziziphus jujuba cv. Dongzao | Rosales | Rhamnaceae | 1.1 | 10.1038/ncomms6315 |
| Zosmar | Zostera marina | Alismatales | Zosteraceae | 2.2 | 10.1038/nature16548 |

**Table S2: Constructs used in this study.** The nomenclature is according to Binder et al., 2014.

| **Golden Gate level** | **DNA-constructs** | **Backbone** |
|---|---|---|
| LII | *pUbi::DREPP-GFP* | BB20 |
| LII | *pUbi::GFP-DREPP* | BB25 |
| LII | *pUbi::DREPP* $^{1\text{-}23}$*-GFP* | BB25 |
| LII | *pUbi::DREPP* $^{24\text{-}214}$*-GFP* | BB25 |
| LII | *pUbi::DREPP*$^{G2A}$*-GFP* | BB25 |
| LII | *pUbi::DREPP*$^{K14E}$*-GFP* | BB20 |
| LII | *pUbi::DREPP*$^{K18E}$*-GFP* | BB20 |
| LII | *pUbi::DREPP*$^{K14/18E}$*-GFP* | BB20 |
| LII | *pUbi::SPR2-mCherry* | BB25 |
| LII | *pUbi::KTN1-mCherry* | BB25 |
| LII | *pUbi::GIP1a-mCherry* | BB25 |
| LII | *pUbi::GCP2-mCherry* | BB25 |
| LII | *pUbi::EB1b-mCherry* | BB25 |
| LII | *pUbi::mCherry-CLASP* | BB25 |
| LIII | *pDREPP::GUS//pUbi::NLS-2xmCherry* | BB53 |
| LIII | *pDREPP::DREPP-GFP//pUbi::NLS-2xmCherry* | BB53 |
| LIII | *pUbi::DREPP-GFP//pUbi::mCherry-SYMREM1* | BB53 |
| LIII | *pUbi::DREPP-GFP//pUbi:FLOT4-mCherry* | BB53 |
| LIII | *pUbi:DREPP-mCherry//pUbi::MAP4-YFP* | BB53 |
| GataWay | *His-DREPP* | pDest17 |

**Table S3: List of primers used for qRT-PCR, genotyping and aomplification of sequences for cloning purposes.**

| Primer for qRT-PCR | Sequence |
|---|---|
| Ubiquitin-f | AAAGCCGGTATAACCGACGATT |
| Ubiquitin-r | TCCAACCCCTCAGAACAACCT |
| DREPP -f | CTGCCGAGAAACAGGAGGAA |
| DREPP -r | CATGCATCAAACTAAGCAAAGCA (in 3´UTR) |
| FLOT4intron f | TAGCTAGCAACAAGATCAATAG |
| FLOT4intron r | AGGGACAGAGTTTAAAGCTC |
| **Primer for genotyping** | **Sequence** |
| Tnt1-F1 | TCCTTGTTGGATTGGTAGCCAACTTTGTTG |
| Tnt1-R2 | CAACAAAGTTGGCTACCAATCCAACAAGGA |
| DREPP-R | AGCCTTTGGTGGTTCTACTTG |
| **Primer for amplification** | **Sequence** |
| DREPP-F | ATGGTCTCACACCATGGGTTACTGGACAACAAAG |
| DREPP-R | TAGGTCTCTCCTTAGCCTTTGGTGGTTCTACTTG |
| pDREPP-F | ATGAAGACTTTACGGGTCTCAGCGGGTGATATTTAGTGAGGGAAAACATG |
| pDREPP-F | ATGAAGACTTCAGAGGTCTCTCAGACTTTTGCTTCCAAATGTTTTTTTTT CTTC |
| $DREPP^{1-23}$-R | TAGGTCTCTCCTTTTTCTTAGTACTATTCTTCTC |
| $DREPP^{24-214}$-F | ATGGTCTCACACCATGACAGCTGCTGCTGAGGTTTCC |
| $DREPP^{G2A}$-F | ATGGTCTCACACCATGGccTACTGGACAACAAAGGTTC |
| DREPP-his-R | TAGGTCTCTCCTTTCAAGCCTTTGGTGGTTCTAC |
| CLASP-F | ATGGTCTCACACCATGGAGGAAGCGTTGGAACTCACAC |
| CLASP-R | TAGGTCTCTCCTTCTAATCATGAACGATATCAATTGC |
| SPR2-F | ATGGTCTCACACCATGAAGAACACAAAACCCCAAACCC |
| SPR2-R | TAGGTCTCTCCTTTTTGTCATGCTGTTGTAGATCAATTTCCC |
| EB1b-F | ATGGTCTCACACCATGGCGACGAACATTGGAATAATGGACG |
| EB1b-R | TAGGTCTCTCCTTTTATGCTGATCCAAATTCATGAAACC |
| KTN1-F | ATGGTCTCACACCATGGCCGGTGGTGGTGGTGGTGG |
| KTN1-R | TAGGTCTCTCCTTTTATGCTGATCCAAATTCATGAAACC |
| GIP1a-F | ATGGTCTCACACCATGGTAACTTTTCTTACTCAAAG |
| GIP1a-R | TAGGTCTCTCCTTAGAAGGAGGTGTAGGATGCAATG |
| GCP2-F | ATGGTCTCACACCATGCCTCGAGAGATAATCACACTTCAGG |
| GCP2-R | TAGGTCTCTCCTTCACTACTGATTTTGGATCCAGTG |

**Table S4: Identifiers for genes used in this study**. Genetic information was used from the Phytozome 12 (https://phytozome.jgi.doe.gov/pz/portal.html) or the *M. truncatula* Genome Database (http://www.medicagogenome.org/).

| **gene name** | **Gene ID** |
|---|---|
| *DREPP* | Medtr2g437530 |
| *CLASP* | Medtr7g099480 |
| *EB1b* | Medtr5g016520 |
| *SPR2* | Medtr3g050060 |
| *KTN1* | Medtr4g011810 |
| *GIP1a* | Medtr2g008230 |
| *GCP2* | Medtr7g117050 |
| *SYMREM1* | Medtr8g097320 |
| *FLOT4* | Medtr3g106430 |
